# Tracking Early Mammalian Organogenesis – Prediction and Validation of Differentiation Trajectories at Whole Organism Scale

**DOI:** 10.1101/2023.03.17.532833

**Authors:** Ivan Imaz-Rosshandler, Christina Rode, Carolina Guibentif, Mai-Linh N. Ton, Parashar Dhapola, Daniel Keitley, Ricard Argelaguet, Fernando J. Calero-Nieto, Jennifer Nichols, John C. Marioni, Marella F.T.R. de Bruijn, Berthold Göttgens

## Abstract

Early organogenesis represents a key step in animal development, where pluripotent cells divide and diversify to initiate formation of all major organs. Here, we used scRNA-Seq to profile over 300,000 single cell transcriptomes sampled in 6 hour intervals from mouse embryos between E8.5 and E9.5. Combining this dataset with our previous E6.5 to E8.5 atlas resulted in a densely sampled time course of over 400,000 cells from early gastrulation to organogenesis. Computational lineage reconstruction at full organismal scale identified complex waves of blood and endothelial development, including a new molecular programme for somite-derived endothelium. To assess developmental fates across the primitive streak, we dissected the E7.5 primitive streak into four adjacent regions, performed scRNA-Seq and predicted cell fates computationally. We next defined early developmental state/fate relationships experimentally by a combination of orthotopic grafting, microscopic analysis of graft contribution as well as scRNA-Seq to transcriptionally determine cell fates of the grafted primitive streak regions after 24h of *in vitro* embryo culture. Experimentally determined fate outcomes were in good agreement with the fates predicted computationally, thus demonstrating how classical grafting experiments can be revisited to establish high-resolution cell state/fate relationships. Such interdisciplinary approaches will benefit future studies in both developmental biology as well as guide the *in vitro* production of cells for organ regeneration and repair.

## Introduction

Single cell transcriptomics has significantly contributed to our understanding of cell type diversity across species, organs and developmental processes. Efforts to build cell atlases from different model organisms include extensive transcriptomic profiling of embryonic development, with profiling of the mouse being particularly relevant given its broad use as a model for mammalian development. Independent efforts now provide coverage from embryonic days E3.5-E6.5, E6.5-E7.5, E4.5-E7.5, E6.5- E8.5, E6.5-E8.25, E9.5-13.5 and E10.5-E15.0 [Mohammed et al. 2017], [Scialdone et al. 2017], [Ibarra-Soria et al., 2018], [Lescroart et al., 2018], [Argelaguet et al., 2019], [Pijuan et al., 2019], [Chan et al., 2019], [Grosswendt et al., 2020], [Mittnenzweig et al., 2021], [Cao et al., 2019], [Han et al., 2018] and [He et al., 2020], complemented by detailed analysis of specific organs such as the brain [La Manno et al, 2021] and the heart [Soysa et al., 2019] or with emphasis on specific germ layers [Nowotschin et al., 2019]. Combining datasets to provide an integrated timecourse over a longer timespan has been achieved by overcoming the challenge of integrating different sequencing technologies [Chengxiang Qiu et al., 2022]. Of note, single cell atlases of normal development have rapidly been utilised as a reference to interpret mutant phenotypes, providing new insights into the cellular and molecular processes controlled by key developmental regulators [Montague et al., 2018], [Pijuan-Sala et al. 2019], [Grosswendt et al., 2020], [Guibentif et al. 2021], [Mittnenzweig et al., 2021], [Barile et al., 2021] and [Clark et al., 2021].

Integrating and annotating transcriptomic profiles in different contexts (e.g., across species, technologies, experiments, time points, etc.) represents a foundational step towards the construction and leveraging of cell atlases, but poses substantial challenges due to the difficulty of distinguishing between highly similar or transitional cell populations arising along complex differentiation trajectories. Without lineage tracing, computational inference of differentiation trajectories may provide useful information for understanding the dynamics of cellular diversification, but need to be interpreted with caution. The increasingly large number of methods for trajectory reconstruction furthermore highlights the need for benchmarking strategies [Sealens et al., 2019]. Many trajectory inference methods are based on the concept of pseudotime, a latent dimension representing the transition between progenitors and differentiating cells as a function of transcriptional similarity. Though related, pseudotime and experimental time are different concepts. Thus, incorporating real time should improve reconstruction of developmental processes [Tritschler et al., 2019]. Inspired by the Waddington landscape [Waddington, 1957] and Optimal Transport theory [Monge, 1781], the probabilistic framework Waddington-OT takes advantage of experimental time and stochastic modelling to estimate the coupling probabilities of cells between consecutive time points and to reconstruct differentiation trajectories [Schiebinger et al., 2019].

Here we report a densely-sampled scRNA-Seq atlas covering mouse development from E6.5 to E9.5 in 6h intervals. This new atlas includes our 116,000 previously published E6.5-E8.5 transcriptomes that cover gastrulation and the initial phase of early organogenesis, complemented by 314,000 new E8.5- E9.5 transcriptomes that bridge a critical gap in development not captured in existing datasets, i.e., the period of major morphological and organogenesis changes that occur between E8.5 and E9.5. This includes embryo turning, emergence of definitive-type Haematopoietic cells, and initiation of the heartbeat and circulation. This new combined E6.5 to E9.5 atlas delivers the most comprehensive transcriptome dataset for mammalian gastrulation and early organogenesis to date. To address the challenge of reconstructing and annotating cell lineages, we combined expert curation with a variety of computational methodologies to generate a resource of broad utility for the developmental biology community. We investigated further the intricate process of haemato-endothelial development, highlighting the multiple origins of endothelial cells and the formation of blood progenitors in asynchronous waves in spatially distinct sites. Moving beyond atlas generation, we also utilised precise embryo dissections to profile spatially-defined areas of the developing embryo. Finally, computational cell fate predictions were contrasted with cell fates observed in state-of-the-art cell grafting experiments, where individual E7.5 primitive streak segments were orthotopically grafted into recipient embryos and resulting cell fates analysed after 24 hours of culture by spatial and scRNA-Seq analysis. Our study provides a blueprint for cell state/cell fate analysis during key stages of mammalian development, paving the way for future interdisciplinary studies of cell fates and origins.

## Results

### A densely sampled scRNA-Seq atlas from E6.5 to E9.5 of mouse development

We previously reported a scRNA-Seq atlas covering mouse gastrulation and the early initiation of organogenesis between E6.5 to E8.5 [Pijuan et al., 2019]. To capture the critical organogenesis period between E8.5 and E9.5 we undertook a new time-course experiment and integrated the new sampling time points (Fig.1a). Thus, 116,312 cells distributed across 9 time points from the original atlas were complemented with 314,027 new cells distributed across four new time points (E8.75-E9.5) as well as one overlapping time point (E8.5) to facilitate data integration (Fig.1b). Combined, the new extended atlas, ranging from E6.5 to E9.5, contains 430,339 cells across 13 time points spanning 3 days of mouse development (Fig.1a-c).

**Figure 1.**
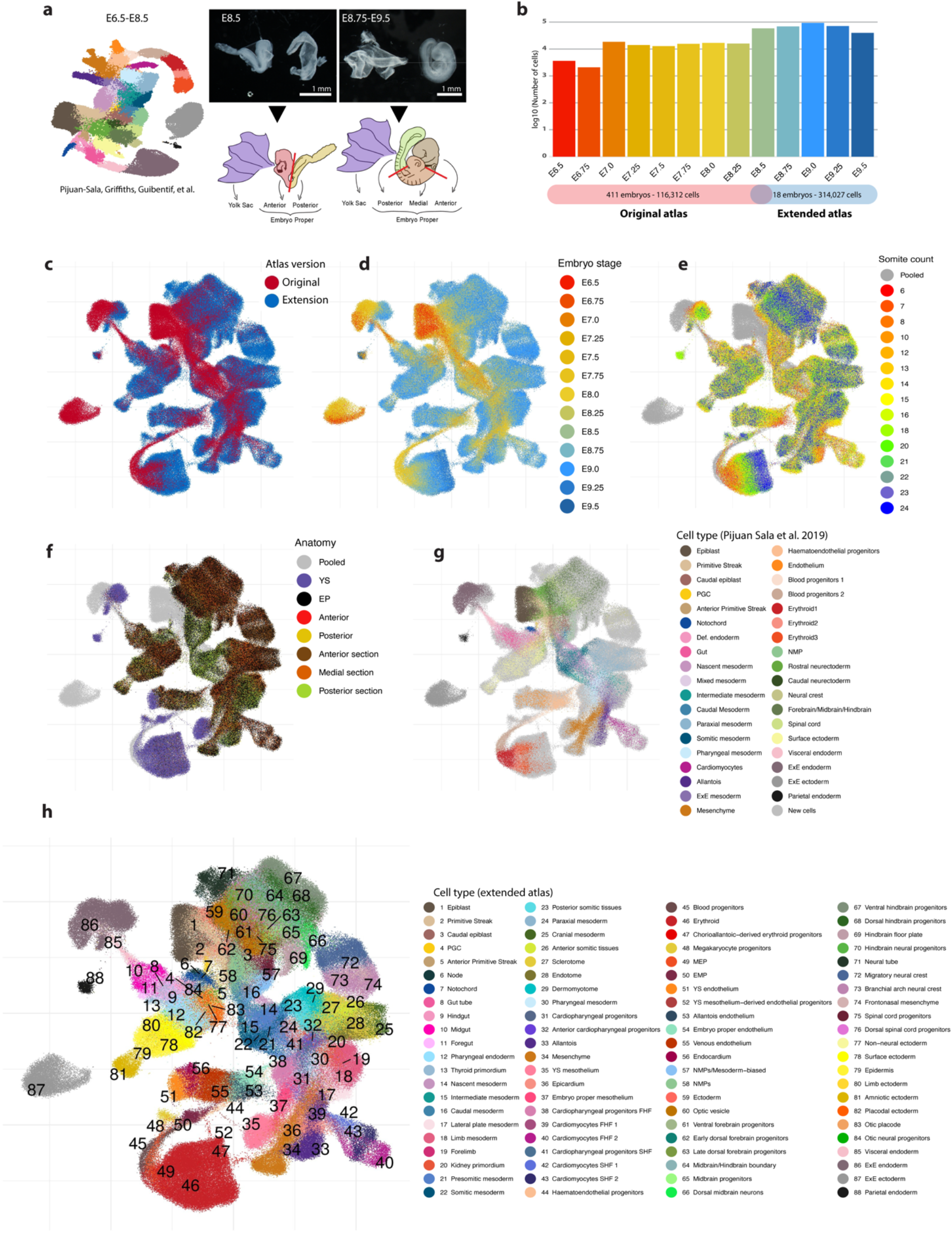
An extended single cell transcriptomic atlas of mouse gastrulation and early organogenesis. **a**) Schematic representation of the experimental design. The publicly available E6.5-E8.5 time course experiment [Pijuan-Sala et al. 2019] was extended towards E9.5 across 24 hours at each six hour interval. An overlapping time point was generated (E8.5) to facilitate batch correction. Information of embryo dissections was recorded to support cell type annotations. **b)** Bar plot showing the number of cells per time point after data integration. In total, the number of transcriptomic profiles increased from 116,312 to 314,027. **c-h)** UMAP layout of the extended atlas. Cells are coloured by **c)** atlas version, the original atlas and the atlas extension, **d)** time-point, **e)** somite counts, as an alternative indication of developmental stage, **f)** anatomical dissection (pooled cells correspond to the original atlas), **g)** cell type annotations provided for the original atlas (newly generated profiles are highlighted in light grey), **h)** cell type annotations resulting from the integration of both atlases and re-annotation process.

All embryos for the new dataset were dissected prior to droplet capture, to (i) profile yolk sac (YS) cells separately and (ii) dissect the embryo proper to provide single cell data anchored by anatomical location (Fig.1a). As illustrated below, sequencing the various tissue segments independently aids the disambiguation of transcriptionally similar cell states (Fig.1d-f). A combination of computational approaches and manual curation (see Methods and Fig.S1) allowed us to define 88 major cell states, more than double the number identified in the previous atlas and reflecting the rapid diversification of cell states during the 24h time window from E8.5 to E9.5 (Fig.1g-h). The new combined dataset has been made freely available through a user-friendly web portal to be explored and leveraged by the wider scientific community (see Data availability).

### Inference of haemato-endothelial development reveals independent intra-and extra-embryonic trajectories

The computational reconstruction of developmental processes using single cell transcriptomics remains a major challenge despite the large number of available methods. The time-course experimental design of this atlas provides the advantage of incorporating developmental stage information thus permitting the use of methods such as Waddington-Optimal Transport (W-OT) that anchor inferred developmental progression in real time rather than pseudotime [Schiebinger et al., 2019]. In W-OT, developmental processes are modelled using a probabilistic framework that allows inferring ancestor and descendant probabilities between cells sampled at consecutive time points, thus allowing for time series analysis. We applied W-OT to the developing haematopoietic and endothelial systems, which are needed early during development to enable effective circulation once the heart starts beating around E8.25-E8.5. Blood and endothelium arise in waves, at least some of which are thought to entail shared haematoendothelial progenitors (reviewed in de Bruijn and Dzierzak, 2017 and Elsaid et al., 2020). The first blood cells are so-called primitive erythrocytes, arising at E7.5 from mesodermal thickenings in the prospective blood islands of the YS. Next is a wave of definitive-type blood progenitors that originate from the YS vasculature from E8.5. The haematopoietic stem cell (HSC) lineage is the last to emerge from the endothelium of major arteries of the embryo, the dorsal aorta and vitelline and umbilical arteries, starting from E9.5. Both YS definitive-type blood progenitors and HSCs are derived from a specialised subset of endothelium, the haemogenic endothelium, through a so-called endothelial-to-haematopoietic transition. The extended mouse atlas covers the two YS-derived waves of blood emergence, as well as a variety of extra and intra-embryonic populations of endothelial cells, providing a unique opportunity to explore the highly complex emergence and diversification of the initial haemato-endothelial landscape during mammalian development.

To encompass all haemato-endothelial lineages, a W-OT fate matrix was computed for all blood and endothelial cell populations in the landscape (erythroid, megakaryocyte, megakaryocyte-erythroid progenitors (MEP), erythroid-myeloid progenitors (EMP), blood progenitors (BP), haemato-endothelial progenitors (HEP), endothelial populations from YS (YS EC), embryo proper (EP EC), and allantois (allantois EC), as well as venous endothelium and endocardium) leaving all other cells in the extended atlas grouped as “other fate” ( Fig.2a). In brief, a W-OT fate matrix is a transition probability matrix from all cells to a number of target cell sets at a given time point, commonly the end point of the time course experiment (here E9.25 and E9.5). Following this strategy, three main W-OT studies were performed. As a first proof-of-concept, only YS blood and endothelial cells were considered with E9.25-E9.5 cells from these two populations defined as targets (Fig.2), which recovered the known divergence between the primitive and definitive-type YS waves over time. Specifically, the W-OT inferred primitive wave mainly generates nucleated primitive erythrocytes, as well as a small number of macrophage and megakaryocyte progenitors (red trajectory in Fig.2b)[Tober et al., 2007]. The second, definitive-type YS wave starts with the emergence of EMPs followed by MEPs [McGrath et al., 2015] arising from YS HE as highlighted by a collection of known markers for all these populations (blue trajectory in Fig. 2b, and gene marker inspection in Fig.2c and Fig.S2, 3 and 4). Lymphoid and microglial-like progenitors can also be detected albeit at low frequency (Fig.S5).

**Figure 2.**
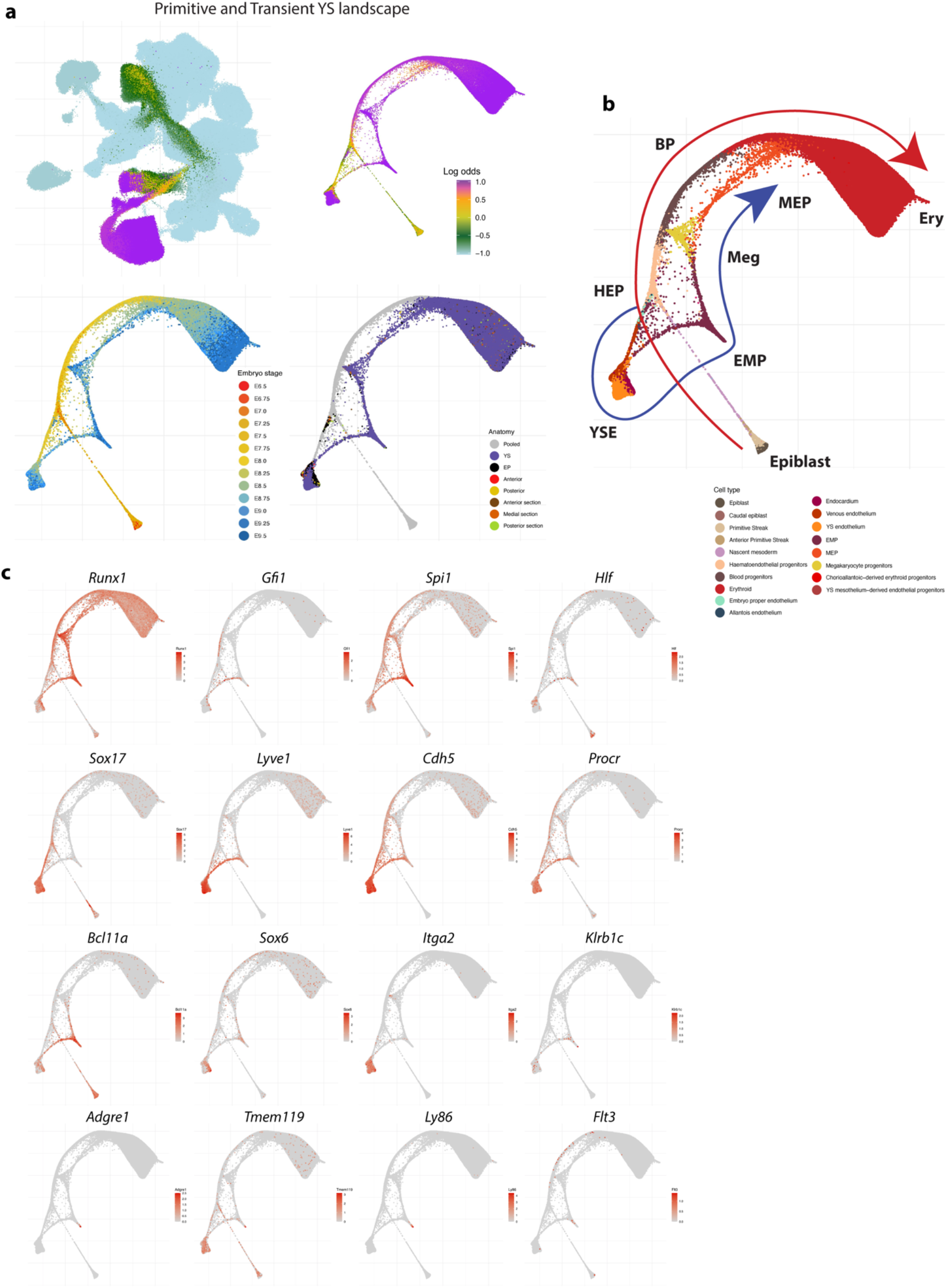
Primitive and definitive YS waves of blood production. **a)** UMAP layout of the mouse extended atlas displaying the log odds of fate probabilities associated with the primitive and YS definitive haemato-endothelial landscape (top left). Cells with log odds > - 0.5 were retained to generate a force directed layout. Cells are coloured by Log odds of fate probabilities of YS blood progenitors and YS endothelial cells, embryo stage and anatomical region. **b)** Force directed layout with cells coloured by cell type, displaying the trajectories of the two distinct waves, clearly distinguished by time. HEP: haemato-endothelial progenitors, BP: blood progenitors. ERY: Erythroid. YSE: Yolk Sack endothelium. EMP: Erythroid-Myeloid progenitors. MEP: Megakaryocyte-Erythroid progenitors. **c)** Force directed layout showing a collection of gene markers associated with these populations, including those represented in small fractions as lymphocytes and microglial progenitors.

For the second W-OT analysis, we wanted to explore the potential origins of endothelial cells together with blood cells, and thus considered the complete haemato-endothelial progenitor population, which resulted in a considerably more complex landscape as endothelial cells are found in multiple regions across the embryo (Fig.3a-b and Fig.S6). Furthermore, we merged blood cells with ‘other fates’ in the W-OT matrix, enabling us to focus explicitly on the endothelial landscape (third W-OT analysis). The endothelial landscape revealed at least three different trajectories (Fig.S7a-b). Differential expression analysis between these populations was performed to identify distinctions between known endothelial markers (Fig.S7c). As expected, YS endothelium expressed *Lyve1*. Embryo proper endothelium was relatively immature as indicated by lower levels of endothelial genes such as *Cdh5*, *Pecam1*, *Dlk1,* as well as the non-coding RNA *Meg3*. Venous endothelium showed the highest expression of genes associated with vasculogenesis and angiogenesis such as *Mef2c, Clec1b and Cldn5*, while the endocardium expressed markers indicative of Bmp signalling (*Id1* and *Id3*) and Notch signalling (*Hey*). In addition to revealing multiple endothelial trajectories, this analysis exhibited a strong connection between somitic tissues and EP ECs as highlighted by the differences in the UMAP coloured by log odds between the first and the second landscape (Fig.2a and Fig.3a). This connection led to the identification of the endotome, a cell population not previously reported in mammalian embryos, which will be further explored in the following section.

**Figure 3.**
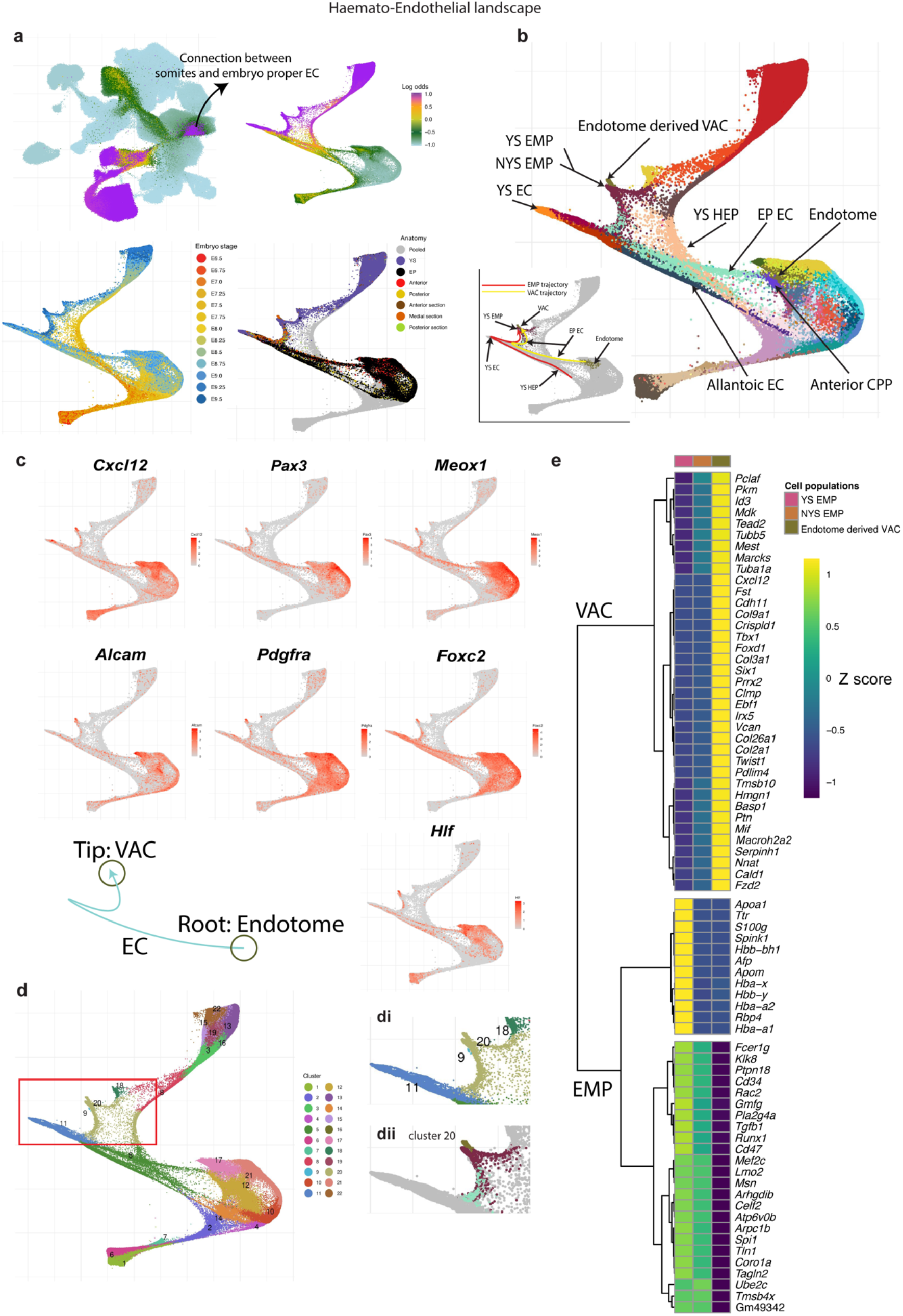
The haemato-endothelial landscape. **a)** UMAP layout of the mouse extended atlas displaying the log odds of fate probabilities associated with the primitive and YS definitive haemato-endothelial landscape (top left). Cells with log odds > - 1 were retained to generate a force directed layout. Cells are coloured by Log odds of fate probabilities of YS blood progenitors and YS endothelial cells, embryo stage and anatomical region. **b)** Force directed layout with cells coloured by cell type, highlighting multiple anatomical origins of haemato-endothelial cells and presence of blood cell types across YS and Embryo proper tissues. A subcluster of Endotome cells is splitted from its major cell type origin and cluster together with EMPs (here named as Endotome derived VACs). Cells are coloured by cell type (see cell type labels at Fig.1h). The bottom left plot further highlights two trajectories and intermediate populations, one for YS EMPs and one for VACs **c)** Collection of gene markers associated with the Endotome population as reported by [Dang Nguyen et al., 2014], [Tani et al., 2020] and *hlf,* which has been associated to HSCs induction [Yokomizo et al., 2019]. Following the landscape in b), the two circles highlight the location of endotome cells and VACs in the landscape, with the trajectory through embryonic endothelial cells indicated by an arrow. **d)** Newly calculated Louvain clusters identified in the landscape. The region of interest is highlighted in the red box. Cluster 20 is extracted and further splitted into anatomically distinct populations and used for differential expression and correlation analysis. Notice that cluster 9 are lymphocyte progenitors. **e)** Heat map displaying differentially expressed genes across different cell populations clustered together in the region highlighted in Dii. YS EMP: Yolk Sack EMP, NYS EMP: Non-Yolk Sack EMP, Endotome derived VAC: Endotome derived vascular associated cells. Mean gene expression values were computed and scaled by rows (Z-score).

### A previously unrecognised developmental trajectory involving novel intraembryonic endotome-like cells

A small but clear subset of endothelium appeared to be part of a continuous differentiation trajectory originating from cells transcriptionally similar to endotome, a somitic subset defined in zebrafish embryos [Dang Nguyen et al., 2014] (Fig.3b). Added credence is given to this transcriptionally defined trajectory because both the contributing endotome and endothelial cells are derived from the same portions of the embryo (anterior/medial regions), demonstrating the utility of subdissecting embryos before generating cell suspensions for sequencing library preparation. Furthermore, consistent with their recent discovery in zebrafish, the mouse endotome cells are characterised by the expression of *Cxcl12, Pax3, Meox1*, *Foxc2*, *Pdgfra* and *Alcam* [Dang Nguyen et al., 2014], [Tani et al., 2020], [Murayama et al., 2023] as well as by *Hlf* which has been associated with intra-aortic haematopoietic clusters and fetal liver HSC induction [Yokomizo et al., 2019] (Fig.3c).

In the zebrafish embryo, endotome cells migrate towards the dorsal aorta and differentiate into endothelial cells that contribute to the niche for the emerging HSCs [Dang Nguyen et al., 2014]. Similarly, earlier studies on chick embryos showed a somite-derived cell population that contributes towards the non-hemogenic endothelium, replenishing vascular cells that have undergone an endothelial-to-haematopoietic transition (EHT) [Pardanaud et al., 1996], [Pouget et al., 2006] and [Sato et al., 2008]. More recently, angioblast-like cells with similarities to somitic mesoderm-derived angioblast and endotome cells of chicken and zebrafish embryos were also reported in pluripotent stem cell-derived human axioloids [Yamanaka et al., 2022]. In lower vertebrate models, the endotome was shown to originate at the ventral–posterior region of the sclerotome [Tani et al., 2020] which is in agreement with the transcriptional neighbours for the new cell population discovered in the mouse embryos here. It took our densely-sampled scRNA-Seq approach to discover the likely connection between the newly-discovered mouse endotome and intraembryonic endothelium (Fig. 3a-c).

Intriguingly, the intraembryonic endothelium was connected with a further, downstream transcriptional state, which was placed in the vicinity of EMP blood progenitors by Force Atlas representations as well as when re-clustering the inferred haemato-endothelial landscape (Fig.3b,d), although previous clustering and cell type annotations over the whole embryo suggested a distinct identity (Fig.1h). Transcriptional analysis revealed this to be a cell state not previously reported in single cell atlases, which lacked a clear haemogenic signature. Instead, these cells likely represent so-called vascular associated cells (VACs, Fig.3b-e and Fig.S8), which are thought to be progenitors of vascular mural and connective tissue cells. Our densely sampled and regionally subdissected atlas therefore suggests substantial transcriptional plasticity and developmental potential within mesodermal cells involved in the formation of intraembryonic blood vessels. Future studies will be required to investigate how this plasticity may extend to the specification of intraembryonic HSCs within the vascular niche at E10.5.

### A gradient of heterogeneous molecular states along the anterior-posterior axis of the primitive streak

The complexity of this atlas is both an opportunity and a challenge for computational inference of developmental processes at the whole organism scale. Importantly, the atlas also provides a framework for complementary experiments which can in turn expand the atlas’ impact. Of particular interest to developmental biologists is the question of whether trajectory reconstruction analysis can unveil the timing at which cells start to differentiate towards either one or a set of specific cell fates. While two-dimensional representations of single cell expression data are often taken as a starting point, such representations lend themselves to overinterpretation, thus highlighting the need for experimental validation. We therefore complemented our extended atlas with (i) scRNA-Seq of carefully dissected subregions of the primitive streak at E7.5 and (ii) orthotopic transplantation of the same streak regions into recipient embryos followed by extended *in vitro* embryo culture and both microscopic as well as molecular fate analysis.

Previous primitive streak cell transplant and labelling experiments produced a fate map of cells along the anterior-posterior axis of the gastrulating mouse embryo revealing reproducible region-specific contribution to all the major cell lineages of the embryo (Kinder et al., 1999). To link these data to the extended mouse atlas, we revisited these experiments and dissected the primitive streak of E7.5 (early bud, EB) mouse embryos into four sequential regions labelled A to D (with A being most posterior and D being most anterior), generated single cell suspensions and profiled the regionally dissected cells using a modified Smart-seq2 scRNA-seq protocol providing deep coverage of each cell (Fig.4a, Fig.S9). We mapped these primitive streak cells onto the extended atlas and employed label transfer to assign cell types (Methods). Approximately 27% of the isolated cells mapped to pluripotent cell types such as epiblast or primitive streak, while 46% mapped to mesoderm, 24% to ectoderm and 3% to endoderm lineages (Fig 4b).

**Figure 4.**
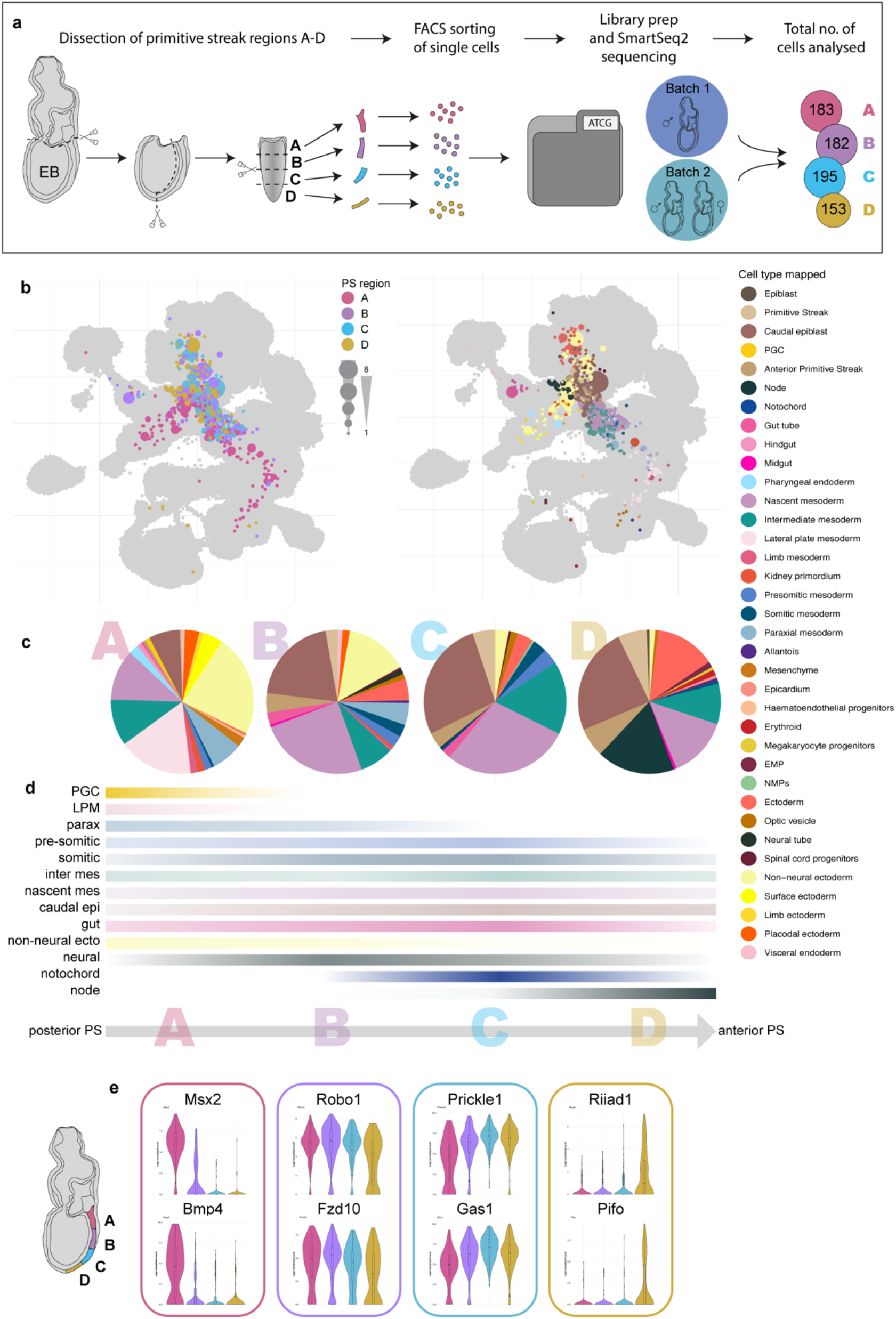
A gradient of transcriptomic differences between four sequential regions of the primitive streak at E7.5. **a)** Schematic of the experimental set up. Single cells isolated from four sequential early bud (EB)-stage primitive streak regions were analysed by scRNA-Seq. A total of 3 primitive streaks were analysed over two experiments. The final cell numbers analyzed for each primitive streak region are indicated. **b)** UMAPs of primitive streak cells mapped onto the extended mouse atlas. Cells are coloured by primitive streak region of origin (left) or transferred cell type label (right). The size of layout dots is proportional to the number of shared closest neighbours across primitive streak cells. **c)** Pie charts showing the relative proportion of the different cell types of individual primitive streak cells mapped to by region of origin (A to D). Five cells from the most distal region D were unexpectedly annotated as erythroid progenitors (Fig.4b, c). These may have inadvertently been included in the analyses during the removal of yolk sac tissues. **d)** Percentile representation of cell type mapping along the primitive streak axis (from A to D, respectively). The colour intensity indicates cell type abundance and resembles the gradient of cell type bias from specific PS portions. **e)** Examples of the most differentially expressed genes between the 4 primitive streak regions.

There were clear differences across primitive streak regions (Fig.4c,e). For instance, only cells from the posterior-most region A mapped to lateral plate mesoderm (LPM) and primordial germ cells (PGCs) as well as expressing known posterior markers such as *Msx2* (expressed in the allantois and known to be involved in PGC migration), and *Bmp4* (Fig 4c,d,e, Fig S10). In contrast, cells from the most anterior region D mapped to the node and expressed several brain and cilia-associated genes, including *Riiad1* and *Pifo* (Fig 4e, Fig S10). Cells from regions B and C were similar to each other and mapped to paraxial, presomitic and somitic mesoderm as well as neural and gut cells. However, these cell types were not unique to regions B and C and were found across the entire posterior-anterior axis of the primitive streak (Fig 4c, d, Fig S10). Consistently, *Robo1*, involved in central nervous system and heart development, *Fzd10*, associated with neural induction, *Prickle1*, a limb development gene, and *Gas1*, a somitic gene also present in LPM, all showed no bias for a particular segment and were expressed throughout the entire length of the primitive streak (Fig 4e). Mesoderm-associated populations such as paraxial, presomitic and somitic mesoderm can be seen as transcriptionally-defined subsets during later development [Guibentif et al., 2020], but these were not yet observed in this EB stage-derived primitive streak dataset. In summary, transcriptional signatures of cells obtained from anterior to posterior primitive streak regions were notably heterogeneous and already showed a bias towards particular molecular states.

### Cell fate analysis of orthotopic primitive streak grafts shows concordance between predicted and observed cells fates

Having identified molecular differences along the anterior to posterior axis of the primitive streak, we next used the extended cell atlas to predict the fates of regions A to D computationally and examined how these compare with experimentally determined cell fates. To this end, using computational fate inference, the closest neighbouring cells of the E7.5 primitive streak cells were identified in the atlas, and downstream fates were inferred based on the W-OT framework outlined above. Based on an initial survey of all predicted fates, a subset of major predictions was selected and visualised in so-called fate plots (Fig.5c and Fig S11e), which revealed associations between predicted cell fates and specific portions of the primitive streak. Cells from the posterior-most region A, for example, were primarily associated with mesodermal fates such as allantois, somites and the non-neural ectoderm, whereas the notochord and neural tube fates clearly favoured the most anterior region D (Fig 5c and Fig S11e).

**Figure 5.**
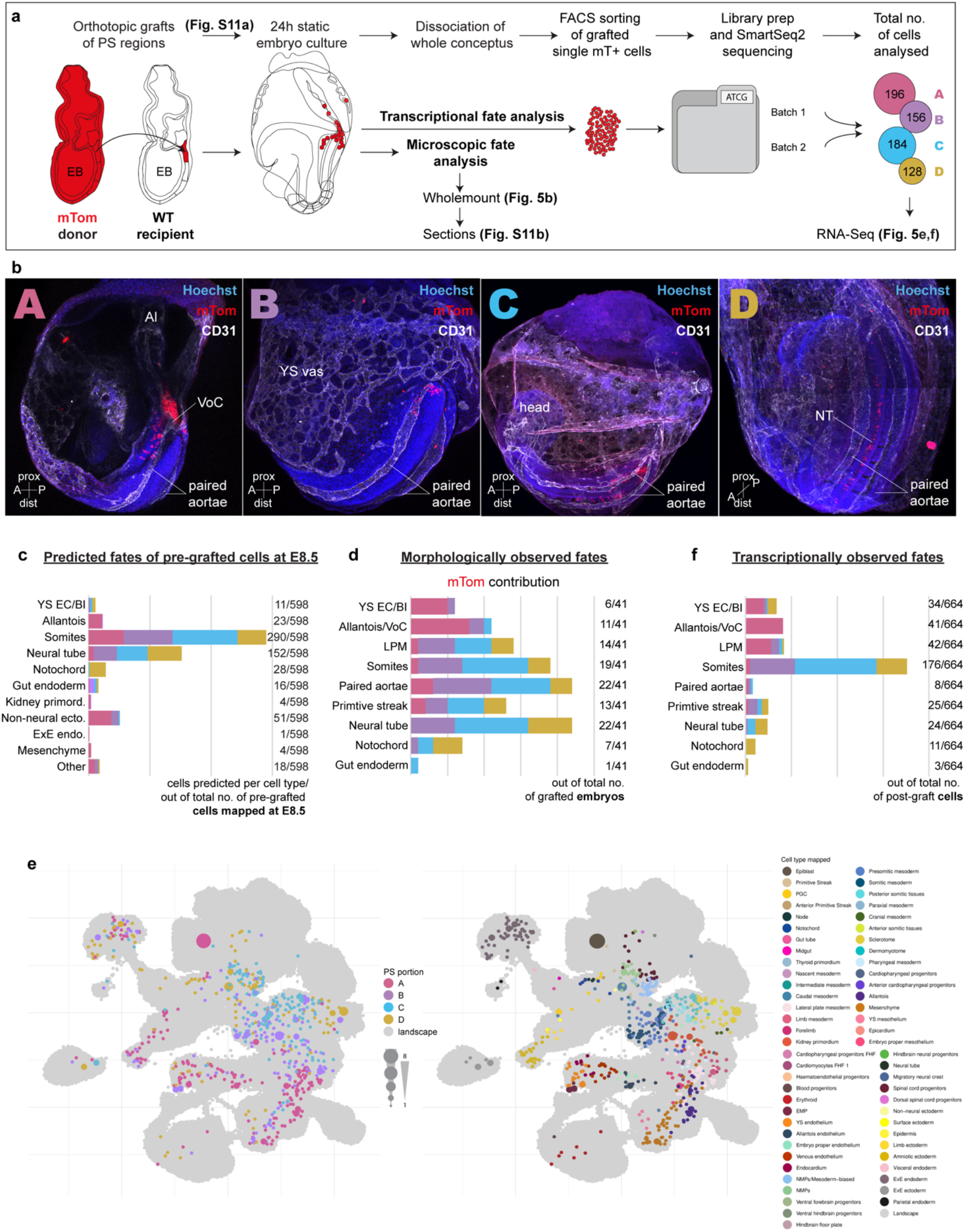
Experimental validation of cell fates in primitive streak grafts. **a)** Schematic of the orthotopic graft experiments with subsequent analysis of cell progeny by imaging and RNA-Seq. Primitive streak regions A to D isolated from E7.5 (EB) mouse embryos carrying a ubiquitous membrane bound tdTomato (mTom) reporter were orthotopically grafted as small cell clusters into E7.5 (EB) wild type recipient embryos. After 24h of culture grafted embryos were either fixed, stained for wholemount analysis and cryo-sectioned for detailed analysis of mTom+ cell contribution; or sorted for mTom+ single cells and processed for SmartSeq2 scRNA-Seq. The total number of mTom+ cells sequenced is given. **b)** Representative wholemount images of grafted embryos after culture, showing mTom (red) contribution to different tissues. Vasculature was stained with CD31 (white). **c)** Fate map of predicted fates of cells freshly isolated from E7.5 (EB) primitive streak regions A to D (documented in Fig.4), determined with the W-OT algorithm. Prediction was capped at E8.5 to allow direct comparison to microscopical and transcriptional fates of the post-grafted donor primitive streak-derived cells. **d)** Fate map of experimentally observed fates of the primitive streak regions A to D based on microscopic analysis (wholemount and sections) of mTom contribution in different tissues of the grafted embryos after culture (Suppl.Table 1). The number of grafted embryos with mTom contribution to particular tissues out of the total number of grafted embryos is shown for each fate. **e)** UMAP of the mouse extended atlas highlighting the closest neighbouring cells to the transcriptomic profiles generated from mTom+ cells isolated from grafted embryos. Cells are coloured by primitive streak region (left) and transferred cell type label (right). The size of layout dots is proportional to the number of shared closest neighbours across primitive streak cells. **f)** Fate map of transcriptionally observed fates based on the mapping of the post-grafted cells on the extended atlas. Only selected cell types are shown here to match the microscopically observed fates. The complete data is provided in Fig. S11d. prox - proximal, dist - distal, A - anterior, P - posterior, Al - Allantois, VoC- Vessel of confluence, YS vas - yolk sac vasculature, NT - neural tube.

To obtain experimental cell fate data, E7.5 embryos were orthotopically grafted with primitive streak regions A to D, cultured for 24h and the fate contributions of the donor regions analysed at E8.25 by microscopic assessment and scRNA-seq (Fig.5a). To facilitate analysis of cell fates, transgenic embryos carrying a ubiquitous membrane-bound tdTomato (mTom, [Muzumdar et al., 2008]) were used as graft donors. Cultured whole embryos and sections were immunostained (Fig 5b, Fig S11b) and careful observation of the location of mTom+ cells showed clear differences in tissue distribution between the donor primitive streak regions (summarised in Fig 5d and detailed in Suppl.Table 1). Notably, region A of the primitive streak contributed predominantly to the most posterior vasculature of the embryo, including the paired dorsal aortae and so-called vessel-of-confluence (VoC, [Rodriguez et al., 2017]), and to the allantois and some YS cells. Regions B and C also contributed to the paired dorsal aortae, as well as to somites, and to the LPM. Region D contributed mostly to the neural tube and notochord (Fig 5d). To analyse the cellular contribution of the grafted primitive streak regions at the transcriptional level, we flow-sorted the mTom+ single cells from additional batches of cultured embryos and performed scRNA-seq (Smart-seq2). After low-level pre-processing, cells were mapped onto the extended atlas as before. Compared to the cells directly isolated from the E7.5 streak, donor primitive streak-derived cells in the E8.25 cultured embryos mapped to almost twice as many cell types (38 vs 63, respectively, compare Fig 4b and Fig 5e), highlighting their advanced differentiation and diversification.

Comparison between the microscopically and transcriptionally-observed fates of the grafted primitive streak regions showed good concordance with only subtle differences in the contribution to distinct lineages (Fig 5d and 5f; microscopically-observed fates are based on embryo counts, while transcriptionally-observed fates are based on individual cell counts). Importantly, comparison between observed and predicted fates showed largely concordant patterns across computationally inferred fates, molecularly (scRNA-Seq) mapped fates and microscopically assigned fates (Fig 5c, d,f). It is worth noting that scRNA-Seq based analysis of fates has substantially enhanced granularity over microscopic analysis, with fate assignment based on our extended atlas for example providing up to 88 different cell types/molecular states for fine-grained annotation. In summary, combining multi-disciplinary approaches, we established primitive streak cell ‘end fates’ at a single cell resolution. Placing these in the context of the extended mouse gastrulation and organogenesis atlas allowed establishing the proof-of-concept for a fate-predictive algorithm, able to forecast fate trajectories of individual primitive streak cells.

## Discussion

To realise the full potential of single cell atlas efforts for developmental biology research, molecular profiling datasets need to (i) provide sufficient sampling density to enable a time-series capable of capturing rapid developmental processes, and (ii) contain sufficient numbers of single cells sequenced at reasonable depth, well annotated and provided to the broader community through user-friendly web portals. Here we report such a resource covering the critical stages of mouse gastrulation and early organogenesis, from E6.5 to E9.5 sampled every 6 hours in 13 individual time steps. This dataset transforms our previous effort by more than tripling the number of cells and more than doubling the number of defined cell states. Furthermore, we revisit classical embryo grafting experiments with transgenic and single cell analysis tools, providing a foundation for future efforts aiming to fully reconstruct cell lineage trees.

Previous single cell atlas efforts from us and others placed observed molecular states into the context of existing knowledge of mouse development (reviewed by [Tam and Ho, 2020] and further integrated by [Chengxiang Qiu et al., 2022]). By contrast, the dense sampling, deep sequencing and regional subdissection of embryos allowed us to discover cell states not previously observed in early mouse development, such as the endotome and VACs. These newly described cell states relate to intraembryonic formation of endothelium, with transcriptomic analysis suggesting that endotome can give rise to endothelial progenitors, which in turn can diversify into VACs already before E9. Similar cell types have previously been described in zebrafish and chicken embryos and recently in pluripotent stem cell-derived human axioloids [Dang Nguyen et al., 2014] [Pardanaud et al., 1996], [Pouget et al., 2006], [Sato et al., 2008], and [Yamanaka et al., 2022], highlighting the value and complementarity of using a variety of different model organisms. Moreover, the zebrafish studies suggested that endotome and VACs may contribute to a cellular niche that promotes intraembryonic formation of blood stem/progenitor cells [Dang Nguyen et al., 2014]. In the mouse, the best-understood site of HSC emergence is the haemogenic endothelium of the dorsal aorta where pro-HSCs are generated from E9.5 [Rybtsov et al.2014]. These haemogenic endothelial cells are lateral plate-/ splanchnopleuric mesoderm-derived and were shown to be replaced by somite-derived endothelial cells concomitant with the extinction of HSC generation [Pouget et al., 2006]. Intriguingly, a recent report suggests that 7 days later during mouse gestation, endothelial cells in fetal bone marrow undergo haemogenic transdifferentiation and produce blood progenitor and differentiated cells [Yvernogeau et al., 2019]. Pax3-Cre lineage tracing further suggested a somitic origin of those haemogenic cells. Future dissection of this process and the underlying mechanisms will be required and will (i) enhance our understanding of early blood and endothelium development, (ii) likely reveal principles relevant to cell plasticity and potential in other developmental contexts, and (iii) provide new mechanistic insights that could be exploited to control differentiation, for example for directed differentiation of pluripotent cells for cell therapy applications.

By combining scRNA-Seq with classical embryo grafting experiments, we show how single cell atlases provide a powerful resource to revisit classical concepts of developmental biology. The results presented in this study are limited in scope but already illustrate how time course-guided trajectory reconstruction performed with computational methods such as W-OT represents a promising approach to dissect complex developmental processes. Our findings also support substantial fate bias present in the E7.5 primitive streak, based on deeply sequenced single cell profiles. Since we did not perform heterotopic transplants, our experiments could not assess fate plasticity nor full fate potential. Moreover, deeper and more comprehensive transplant experiments should be designed around new experimental tools not available when this work was performed, including extended *in vitro* embryo culture [Aguilera-Castrejon et al., 2021], as well as single cell barcoding to permit reconstruction of single cell phylogenies [Bowling et al., 2021].

A future thereby emerges where a confluence of complementary technologies will transform our understanding of early mouse development to a level previously only attained with non-mammalian organisms. Building on data resources such as the one reported here, mechanistic insights will continue to require perturbation experiments, with the important proviso that carefully chosen experimental perturbations have the added benefit to provide new insights into disease processes, in particular congenital defects associated with mutations in developmental regulator genes. Embracing this future is likely an important prerequisite for developmental biology research to remain a prominent component of leading academic research centres.

## Materials and methods

### E8.5-E9.5 embryos collection for the extended atlas

Mouse embryos were collected under the project licence number PPL 70/8406. Animals used in this study were 6-10 week-old females, maintained on a lighting regime of 14h light and 10h darkness with food and water supplied *ad libitum*. Following wildtype C57BL/6 matings, females were killed by cervical dislocation at E8.5, E8.75, E9.0, E9.25 and E9.5. The uteri were collected into PBS with 2% heat-inactivated FCS on ice and the embryos were immediately dissected and processed for scRNA-seq. For each timepoint, four embryos were selected based on morphology and somite counts, in order to span the range expected for the given timepoint according to [Theiler, K., 1989] and processed individually. The exception is the E9.5 time-point, where embryos were smaller than expected and only two embryos were collected with lower somite numbers. The yolk sac was systematically separated from the rest of the embryo and processed as a separate sample. Of the four selected embryos at each time-point, two were partitioned in defined anterior-posterior sections dissociated as separate samples, and two were dissociated as bulk and the suspension then divided into two separate samples for 10X RNA-Seq analysis. For the E8.5 partitioned embryos, they were divided into two halves with the cut being made at the 4th somite level (i.e., right before the 4th somite level). The anterior portion (including headfolds, branchial arches and heart rudiment) and the posterior portion (incl. allantois, hindgut, primitive streak) were processed as individual samples. For E8.5 bulk-dissociated embryos, the single-cell suspension was divided into two separate samples for 10X RNA-Seq analysis.

E8.75-E9.5 embryos were divided into 3 segments, with cuts made below the otic pit and below the heart (at the 10th-12th somite level). The anterior most portion (incl. brain structures anterior to rhombomere 6 and branchial bars), mid-portion (incl. the heart and remains of vitelline vessels) and posterior portion (incl. allantoic structures, hindgut and posterior most somites) were then singularized and further processed as separate samples. Single cell suspensions were prepared by incubating the samples with TrypLE Express dissociation reagent (Life Technologies) at 37 °C for 7 min under agitation and quenching in PBS with 10% heat-inactivated serum. The resulting single-cell suspension was washed and resuspended in PBS with 0.4% BSA and filtered through a Flowmi Tip Strainer with 40 μm porosity (ThermoFisher Scientific, # 136800040). Cell counts were then assessed with a haemocytometer. Single-cell RNA-seq libraries were generated using the 10X Genomics Chromium system (version 3 chemistry), and samples were sequenced according to the manufacturer’s instructions on the Illumina NovaSeq 6000 platform.

### Publicly available mouse gastrulation data

The mouse gastrulation atlas was processed exactly as described in [Pijuan Sala et al., 2019]. This time-course experiment contains 116,312 cells distributed across 9 time points sampled across E6.5-E8.5 at six hours intervals. Only two samples of this dataset (where a sample is a single lane of a 10x Chromium chip) contained pooled embryos staged across several time points. Cells from these samples are denoted as ‘Mixed gastrulation’ in the metadata.

### 10x Genomics data Low level analysis

Raw reads were processed with Cell Ranger 3.1.0 using the mouse reference 1.2.0, mm10 (Ensembl 92) and default mapping arguments. The following steps of pre-processing were performed with R using the same functions, parameters and software versions broadly described in [Pijuan Sala et al., 2019]: swapped molecule removal, cell calling, quality control, normalisation, selection of highly variable genes, doublet removal, batch correction and stripped nucleus removal. Therefore, singularity containers available in https://github.com/MarioniLab/EmbryoTimecourse2018 provide the necessary software for reproducing these steps.

### Generating an integrated atlas

Log transformed normalised counts obtained from [Pijuan Sala et al.] and those generated here were integrated into an extended time course experiment across mouse developmental stages E6.5-E9.5 (13 time points). Hence, this expression matrix contained 23,972 genes and 430,339 cells. Selection of highly variable genes and batch correction with fastMNN function from scran [Lun et al., 2016] was performed as described above, resulting in 5,665 genes and 75 batch-corrected principal components.

### Mapping stage and cell type annotations within the extended gastrulation atlas

Metadata annotations such as embryonic developmental stages and cell types were assigned to the mixed time points (annotated as Mixed gastrulation in the metadata) and newly generated E8.5 samples respectively using a strategy based on fastMNN. In this approach, UMI counts from both, the reference and the query datasets, are merged, normalised and log transformed together. Then, highly variable genes and top principal components are computed to subsequently use fastMNN for re-scaling the PCA space from both datasets. The annotations from the reference metadata are assigned to the query data as the mode among k nearest neighbours (KNN) between the query and the reference PCA subspaces using queryX function from Biocneighbours. The number of nearest neighbours is chosen depending on the resolution of transferred annotations. Mixed gastrulation time points were allocated to embryonic developmental stages using the 30 nearest neighbours queried from the top 50 batch-corrected principal components from the E6.5-E8.5 reference dataset. New E8.5 cell type annotations were assigned using 10 nearest neighbours with respect to E8.5 cells from the reference atlas in the corresponding E8.5 subspace of the integrated batch corrected PCA described above.

### Constructing optimal transport maps

The W-OT approach was conceived to model time course experiments of developmental processes as a generalisation of a stochastic process using unbalanced optimal transport theory. Thus, it allows estimating the coupling probabilities between cells of consecutive time points, while taking into account cell growth and death [Schiebinger et al., 2019]. The transport maps of consecutive time points were constructed over the entire set of cells with Waddington-OT (wot 1.0.8.post1) using default settings except for skipping the dimension-reduction step, and instead using the batch-corrected principal components as input as well as three iterations for learning the cell growth rate.

Embryonic stages E9.25 and E9.5 were collapsed into a single time point for computing the transport maps because the somite count (a more accurate measure of developmental stage), overlapped between these embryos.

### Estimating the descendants from cell populations at E8.5

The transport maps above explained were used to estimate the full trajectories of every cell population present at E8.5. That is, for each cell population (i.e., Erythroid3) the coupling probabilities were used to reconstruct the sequences of ancestors and descendants distributions by pushing the cell set through the transport matrix backwards and forwards respectively. Cells were allocated as descendants from a cell population at E8.5 by selecting those with maximum mass across all trajectories in time points E8.75-E9.5.

### Integrating the brain and gut development atlases to support cell type annotations

Mapping publicly available data from the atlas of brain development [La Manno et al., 2021], and both atlases of gut development [Nowotschin et al., 2019] against the extended gastrulation atlas was performed following the strategy based on fastMNN above mentioned. However, in this case the matrices of principal components from the query dataset were randomly split into subsets smaller than 10,000 cells for fastMNN and merged back to generate a single mapping output.

### Expansion and refinement of cell population annotations

A combination of complementary strategies was used for defining final cell type annotations (Fig S1). First, cell type annotations from [Pijuan et al., 2019] were transferred within overlapping time points (E8.5) and cell descendants were estimated as described above (Supp. Figure 1a-b). The landscape was then split into two subsets, a mesodermal and an ectodermal-endodermal landscape (the latter including NMPs) (Fig S1c), and subsequently clustered using scanpy’s implementation of Leiden’s algorithm [Tragg et al., 2019] with resolution 5 (Fig S1d). Highly variable genes and batch corrected principal components were recomputed on the subsets before clustering. Then, annotations from the original atlas expanded through mapping and estimation of cell descendants were manually refined by means of differential expression analysis between clusters (using findMarkers from scran’s package version [Lun et al., 2016]), as well as literature and visual inspection of gene markers resulting in major 88 cell type populations (Figure 1h).

### Generating the landscapes of haemato-endothelial trajectories

The strategy for generating the haemato-endothelial landscapes was based on the so-called W-OT fate matrix. In brief, a W-OT fate matrix is a transition probability matrix (the rows of this matrix add up to 1) from all cells to a number of target cell sets at a given time point, commonly the ending point of the time course experiment. To fully cover the haemato-endothelium, three W-OT fates matrices were computed, one for each landscape presented in our results. One using E9.25-E9.5 YS-blood and YS endothelial cells as targets, another one with YS blood as well as both, YS and embryonic endothelial tissues, and a third one where only endothelial cell types were considered targets (Figures 2, 3 and 4 respectively). Then, to identify cells that form a differentiation trajectory towards any of the above mentioned cell fates, for every cell in the landscape the likelihood probabilities associated with these fates and the remaining cells at E9.25-E9.5 cell types (i.e., cells grouped as "Other fate") were compared using the log odds or ratio of probabilities. The log odds is defined as the logarithm of the ratio of probabilities of two different categorical and mutually exclusive outcomes *log*(*p*/1 − *p*). These ratios provided a quantitative setting for more interpretable thresholds when selecting cells that potentially belong to a cell fate trajectory. These cells are then retained and used to generate new layouts including only potentially fated cells towards blood and endothelium. That is, selecting pluripotent and mesodermal cells above a reasonable log odds threshold, recomputing highly variable genes, correcting for batch effects and generating new force directed graphs from inferred trajectories. Cells with log odds > 0 were considered fate biased towards the set of selected population targets, such as in the proof of concept performed in Fig.2. Since the log odds is computed by summing over the cell probabilities for all population targets (here denoted as p), divided by the probability of all other fates grouped together (1-p); cells with negative values but close to 0, although with higher uncertainty might well be contributing to the population targets of interest. Thus, for more complex landscapes the threshold was lowered to log odds > −1 (Fig.3 and Fig.S7), in order to exclude all cells not associated with haematoendothelial fates with high confidence, while keeping a higher degree of uncertainty for those retained.

### Differential gene expression and Canonical Correlation Analysis of Endotome derived VACs

Subsequent to the construction of a complete haemato-endothelial landscape, louvain [Subelj and Bajec et al., 2011] clusters (as implemented in the R package igraph [Gabor and Tamas, 2006]) were generated from the batch corrected principal components obtained as explained above (Fig.3c). Cluster 20 was of particular interest for investigating both shared and non-shared gene expression profiles between the anatomically distinct populations YS EMP and NYS EMP, and a subset of endotome cells that is placed next to them (Fig.3d-e). Differential gene expression analysis was performed using the function findMarkers from the scran package [Lun et al., 2016]. The top 20 ranked genes from each group with a mean log fold change greater than 0.5 were selected for visualisation using a heatmap. CCA as implemented in the R package CCA [Gittins, R. 1985] and [Mardia et al., 1979] was applied to mean metacell expression values of YS EMPs and the endotome derived VACs. Positively correlated genes were then identified by selecting only those with positive coefficients (canonical scores) over the two canonical variates and visualised using a scatter plot fitting a linear regression. Importantly, such analysis was performed using Metacells [Baran et al., 2019] to strengthen gene expression signals.

## Metacells

We identified metacells (i.e., groups of cells that represent singular cell-states from single-cell data) with the goal of achieving a resolution that retains the continuous nature of differentiation trajectories while overcoming the sparsity issues of single-cell data. We adopted the SEACells implementation [Persad et al., 2022]. Following the method guidelines, metacells were computed separately for each sample using approximately one metacell for every seventy-five cells. Following metacell identification, we regenerated the gene expression matrices summarised at the metacell level. Sample-specific count matrices were then concatenated and normalised together. Metacells were used exclusively to inspect the correlation between genes expressed in YS EMPs and endotome derived VACs, as well as the lack of a clear hemogenic signature. However, metacells were computed for all the haematoendothelial landscapes.

### Smart-seq2 data low level analysis

Sequencing reads were aligned against the *mm10* genome following the *GRCm38.95* genome annotation (*Ensembl 95*) using *STAR* version 2.5 [Dobin et al., 2003] with --intronMotif and 2-pass option mapping for each sample separately (i.e., --twopassMode basic) to improve detection of spliced reads mapping to novel junctions. Sequence alignment map (SAM) files were then processed with *samtools-1.6* in order to generate the corresponding count matrix with *htseq-count* from *HTSeq*, version 0.12.4. Cells with total counts lower than 10,000 reads were filtered out from downstream analysis based on observed library size distributions. The mitochondrial fraction of reads per cell and library complexity were computed as part of the QC. Only two cells showed a mitochondrial fraction larger than 4% and the lowest number of genes detected was 3,113. Thus, no cells were excluded during QC. Size factors were computed with *scran* and log transformed normalised counts were obtained with the *logNormCounts* function from *scater* using size factors centred at unity prior to calculation of normalised expression values. Then, highly variable genes were extracted using *ModelVar* function from *scran*, according to a significant deviation above the mean-variance fitted trend (BH-corrected p<0.05). These genes were selected for computing a PCA. The top 50 PCs were retained for batch correction using *reducedMNN*, which is an updated version of *fastMNN*. Figure 4.3 shows the two-dimensional PCA, before and after batch correction. The resulting batch-corrected PCs were used for clustering and visualisation in downstream analysis.

### Predicting transcriptional identity and cell fate of primitive streak cells by mapping them against the atlas

Following the strategy to transfer annotations from the atlas to other datasets previously described above, cell type annotations were assigned to primitive streak cells according to the 15 nearest neighbours between the two manifolds (query data and atlas reference). The annotation transfer was performed separately for each batch to avoid introducing an extra confounding factor when applying MNN batch correction. Also, prior to mapping against the atlas, genes with total counts lower than 100 were filtered out from the Smart-seq2 datasets due to the large differences in dropouts. The resulting annotations were projected onto the UMAP extended atlas embryo landscape by highlighting the closest cells on it. By retaining the closest cell in the atlas and selecting the cell fate with highest probability allocated in a W-OT fates matrix, we aimed to not only map the observed transcriptome at E7.5 but make predictions about likely cell fates at future timepoints (eg. E8.5 and E9.5).

### Primitive streak dissections and processing for Smart-Seq

Mouse embryos were collected under the UK Home Office project licence number PP9552402. Animals used in this study were 6-16 week-old wild type females of a mixed (CBAxC57BL/6)F2 background, maintained on a lighting regime of 14h light and 10h darkness with food and water supplied *ad libitum*. Following timed mating crosses between transgenic mTmG (B6.129(Cg)- *Gt(ROSA)26Sor^tm4(ACTB-tdTomato,-EGFP)Luo^*/J [Muzumdar et al., 2007]) males and wildtype females, females were killed 7 days after a vaginal plug was observed via a Schedule 1 method. The uterine tissue and the decidua were cut away to extract the E7 embryo. The Reichert’s membrane was peeled off and embryos were staged according to [Downs and Daviers, 1993]. Early bud (EB) stage embryos were taken for further analysis. For primitive streak dissections, the extra-embryonic part of the conceptus was cut off, the anterior and posterior embryonic portions were separated, and the posterior portion flattened out, taking care not to mix up the orientation. The posterior portion containing the primitive streak was cut at the halfway point and each half was halved again creating 4 similar-sized primitive streak segments. All embryo and primitive streak dissections were performed in Dulbecco’s Phosphate

Buffered Saline (PBS; with CaCl2 and MgCl2, Gibco) supplemented with 10% FCS (Gibco), 50U/ml penicillin and 50U/ml streptomycin.

For scRNA-seq of primitive streak cells, individual primitive streak regions were collected in separate Eppendorf tubes in FACS buffer (PBS without CaCl2 and MgCl2 (Gibco) supplemented with 10% FCS (Gibco), 50U/ml penicillin and 50U/ml streptomycin). After centrifugation at 200G for 5 min, tissues were resuspended in 200-250ul of TrypLE Express (Life Technologies) and incubated at 37C for 7 min with regular agitation and dissociated by pipetting. Cells were washed with 1ml of FACS buffer, centrifuged at 200G for 5min and resuspended in FACS buffer with Hoechst 33358 for sorting. In order to reduce cell loss from these small cell populations, wild type adult mouse thymocytes were added to the samples and were gated out based on their typical FSC-SSC profiles and absence of tdTomato expression. Each primitive streak region yielded on average 200 cells. Cells were kept on ice throughout the procedure. Individual cells were sorted directly into 96 well plates (Starlab E1403- 1200) into 2.3ul of lysis buffer containing SUPERase-In RNase Inhibitor 20U/ul (Ambion, AM2694), 10% Triton X-100 (Sigma, 93443) and RNAse-free water. Plates were processed using a combination of Smart-Seq2 [Picelli et al., 2014] and mcSCRB-seq [Bagnoli et al., 2018] For detailed protocol see [Sturgess et al., 2021] Pooled libraries were run on the Illumina HiSeq4000 at Cancer Research UK Cambridge Institute Genomics Core.

### Primitive streak grafts and static culture

mTmG transgenic embryos were dissected to harvest 4 equal primitive streak regions as described above. Each segment was deposited into a 50ul dissection buffer drop on the inside lid of a 5cm dish and labelled. One larger drop was deposited in the centre into which up to 5 dissected wild type embryos of the same stage (EB) were transferred. Primitive streak regions were grafted into orthotopic positions in the wild type embryos using a pulled glass capillary needle attached to a mouth pipette. Cells from each donor region were divided over 2-3 recipient embryos with each recipient embryo receiving 50-100 primitive streak cells, mostly as a chain of cells rather than in suspension. Embryos were gently washed in a drop of culture media and gently transferred into individual wells of a 96 well Ultra-Low adhesive plate (Corning) into 200ul culture media. Culture media was 100% rat serum (Envigo, custom collected). Embryos were cultured at 37C, 5% CO2 for 23-26h.

### Fate contribution analysis

Embryos were taken out of the incubator and assessed for (the beginnings of) a heartbeat. Only embryos that lacked deformities and had a heartbeat were used for further analysis by microscopy or scRNA-Seq to assess the fates of the grafted primitive streak cells. For microscopically observed fates, reporter gene expression (mTom) was enhanced (anti-RFP antibody) and embryonic vasculature (CD31) visualised by immunostaining. Wholemount embryos were imaged to assess overall contribution of mTom in relation to embryo anatomy. Embryos were next embedded and cryosectioned, followed by imaging of all areas with mTom+ contribution and careful analysis and scoring for detailed contribution to a selection of most anatomically distinct cell lineages (Suppl Table 1). For scRNA-seq, grafted, post-culture embryos were staged by somite pair counts and individual donor embryo and single cell suspensions prepared for sorting as described above for E7.5 primitive streak region. Single mTom+ cells from the grafted embryos were sorted into 2.3ul lysis buffer in 96 well plates and processed for Smart-Seq2 as above.

## Data availability

This data has been made available at https://marionilab.github.io/ExtendedMouseAtlas/ where links to both, raw and processed data are found. Memory usage to process and analyse this data was optimised by means of *scarf*, an efficient single cell analysis framework [Dhapola et al., 2022].

## Acknowledgements/Disclosure statements

We would like to thank all the people and the different funding instances contributing to this project. Work at Cambridge was supported by Wellcome Collaborative award 220379/B/20/Z (Cambridge Gastrulation Consortium), Wellcome award 221052/C/20/Z (Human Cell Atlas - development atlas extension), and Wellcome award 206328/Z/17/Z (Defining the Haematopoietic System through Integrated Multi-Scale Analysis). Work in the Gottgens group was supported by Wellcome, Bloodwise, MRC, CRUK, by core support grants from Wellcome to the Wellcome-MRC Cambridge Stem Cell Institute. We thank Sarah Kinston for technical assistance. Work in the Marioni group was supported by core funding from CRUK (C9545/A29580) and by the European Molecular Biology Laboratory. J.C.M. has been an employee of Genentech since September 2022. Work at Oxford in de Bruijn group was supported by programmes MC_UU_00016/2 (MdB and CR) and MC_UU_00029/5 (MdB) in the MRC Molecular Haematology Unit Core award. We thank Biomedical Services staff at Oxford for animal care. We would like to acknowledge Kevin Clark in the flow cytometry facility at the MRC WIMM for providing cell sorting services. The facility is supported by the MRC HIU; MRC MHU (MC_UU_12009); NIHR Oxford BRC; Kay Kendall Leukaemia Fund (KKL1057), John Fell Fund (131/030 and 101/517), the EPA fund (CF182 and CF170) and by the MRC WIMM Strategic Alliance awards G0902418 and MC_UU_12025. We thank Christoffer Lagerholm and Jana Koth of the Wolfson Imaging Centre Oxford for their assistance. The centre is supported by the MRC via the WIMM Strategic Alliance (G0902418), the MHU (MC_UU_12009), the HIU (MC_UU_12010), the Wolfson Foundation (18272) and the Wellcome Trust (Micron 107457/Z/15Z) grants. I.I. was funded by the Wellcome Mathematical Genomics and Medicine Programme at the University of Cambridge (203942/Z/16/Z RDAG/426 RG86191). C.G. was funded by the Swedish Research Council (2017-06278) and by a Swedish Childhood Cancer Fund position grant (TJ2021-0009). M.-L.N.T. was funded by a Herchel Smith PhD Fellowship in Science. D.K. was funded by the Wellcome Mathematical Genomics and Medicine Programme at the University of Cambridge (PFZH/158 RG92770).

## Author Contributions

J.N. set up mouse timed mating. C.G., M.-L.N.T. and J.N. performed embryo dissection for the atlas.

C.G. and F.J.C., performed sample and sequencing library preparations for 10X scRNA-Seq. I.I., performed the whole computational analysis except for Metacells. R.A. performed Metacells analysis. I.I, C.G., M.-L.N.T., and D.K. annotated atlas cell types. I.I. and P.D., developed the shiny app with scarf in the backend. C.R. performed Primitive Streak dissections and sample preparation for Smart-Seq2 experiments, Primitive Streak grafts and static cultures. I.I. and C.R. prepared the manuscript figures. J.C.M, M.B. and B.G. supervised the study. I.I., C.R., C.G., J.C.M, M.B. and B.G wrote the manuscript. All authors read and approved the final manuscript.

## Supplementary Figures

**Supplementary Figure 1.**
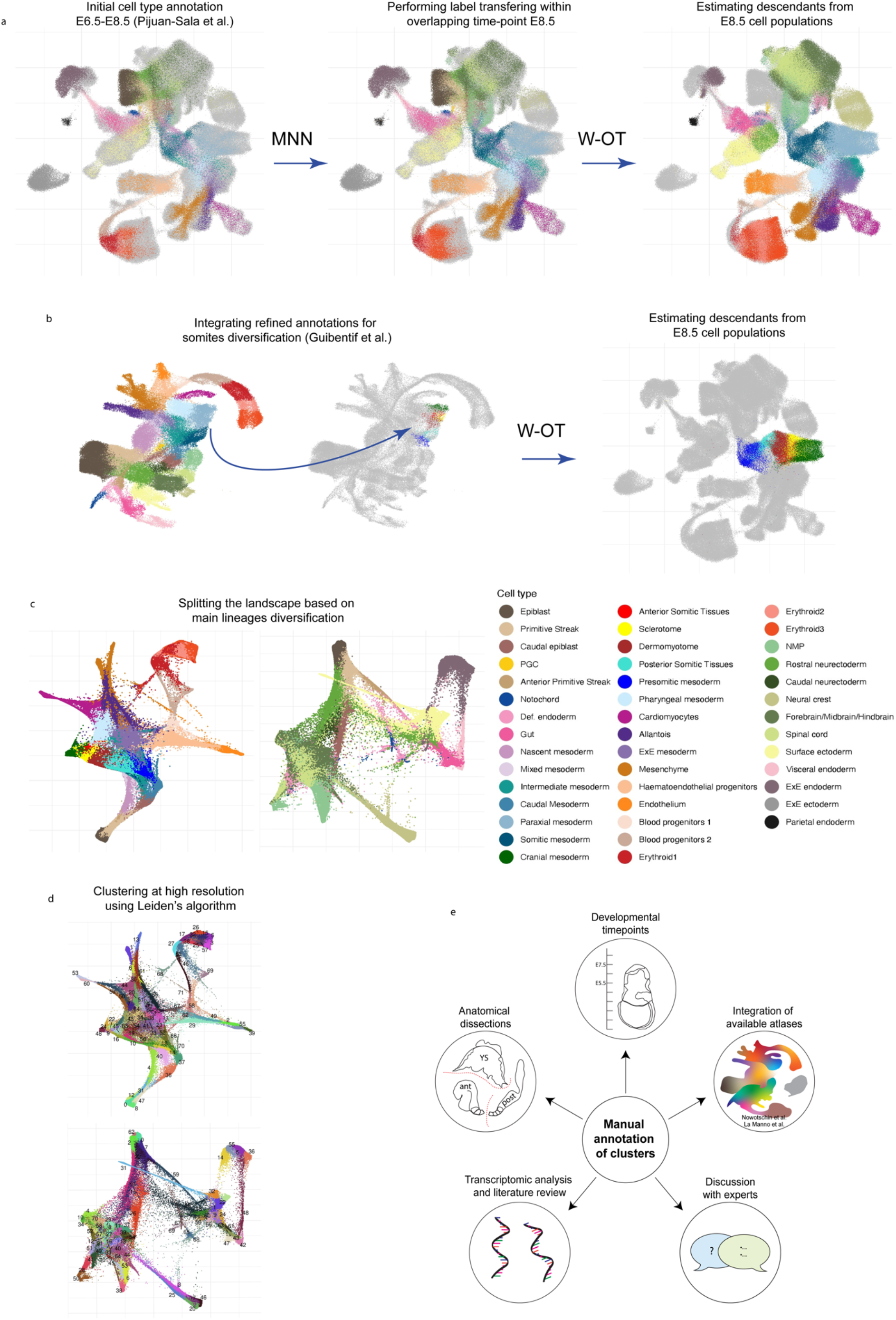
Extending cell type annotations step by step. **a**) UMAP layouts are used to visualise the first stage of cell type annotation, cells are coloured by cell type [Pijuan-Sala et al. 2019] but light grey cells represent those new that have not yet been annotated. Initially, mutual nearest neighbours (MNN) were used to transfer E8.5 cell type annotations from [Pijuan-Sala et al. 2019] to newly profiled cells within the overlapping time-point. Next, cell descendants are estimated for all existing E8.5 cell populations using W-OT by pushing forward the mass over the transport maps in the following time-points (E8.75-E9.5). Here, only cell descendant populations are coloured by cell type and light grey cells refer instead, to those from the original atlas (E6.5-E8.5). **b)** Refined cell type annotations for Paraxial and Somitic mesoderm from Guibentif et al. [Pijuan-Sala et al. 2019] are integrated through the same process described in a). **c)** The atlas is splitted into two landscapes based on the annotations deriving from a) and b). The mesodermal landscape at the left and the ectodermal/endodermal one at the right. Cells are coloured by original Atlas cell types and descendants. **d)** Cells are coloured by high resolution louvain clustering as the starting point of new cell type annotations. **e)** Highly variable genes, batch correction and leiden clustering was performed independently for the two landscapes obtained from C. These clusters are manually annotated using different sources of information as illustrated in the right side scheme. For instance, cell type annotations from other atlases such as [Nowotschin et al. 2019 and La Manno et. al, 2021] were used as guidance, differential expression analysis and literature marker inspection and trajectory analysis.

**Supplementary Figure 2.**
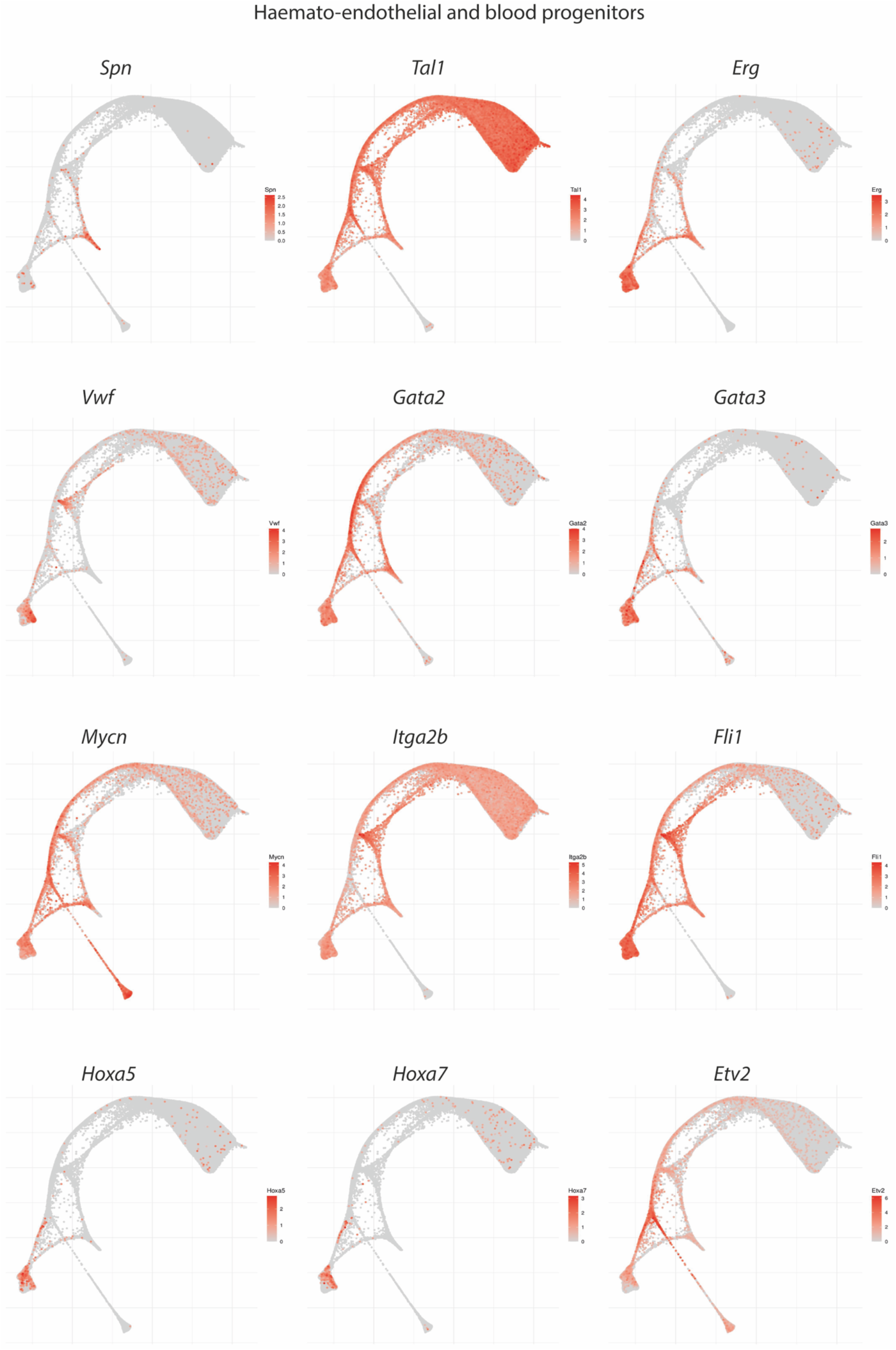
Haemato-endothelial and blood progenitors. Collection of gene markers for haemato-endothelial and blood progenitors shown as a complement for Figure 2. Force directed layouts displaying gene markers expression levels.

**Supplementary Figure 3.**
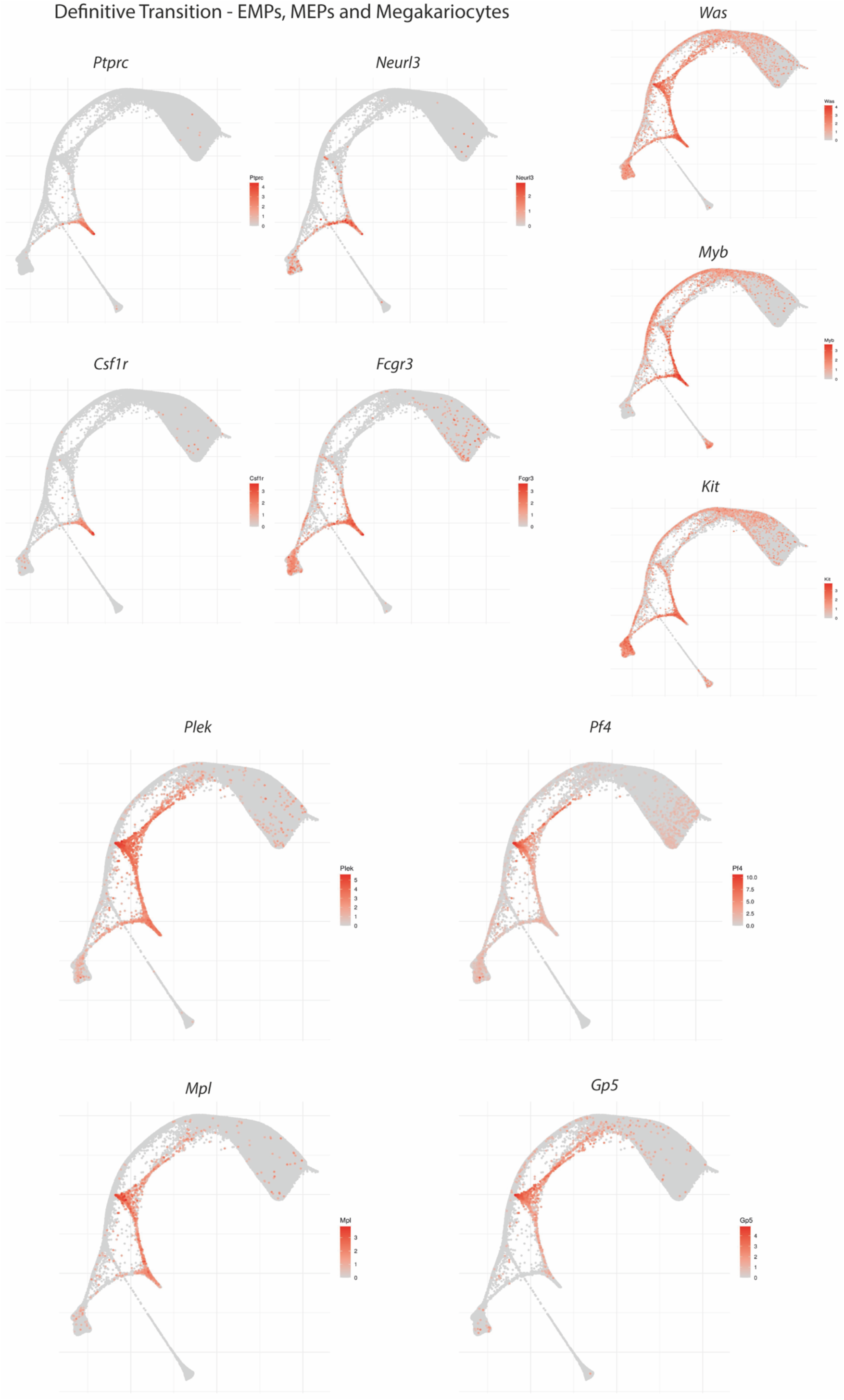
Definitive transition - EMPs, MEPs and megakaryocytes. Collection of gene markers for definitive blood populations shown as a complement for Figure 2. Force directed layouts displaying gene markers expression levels.

**Supplementary Figure 4.**
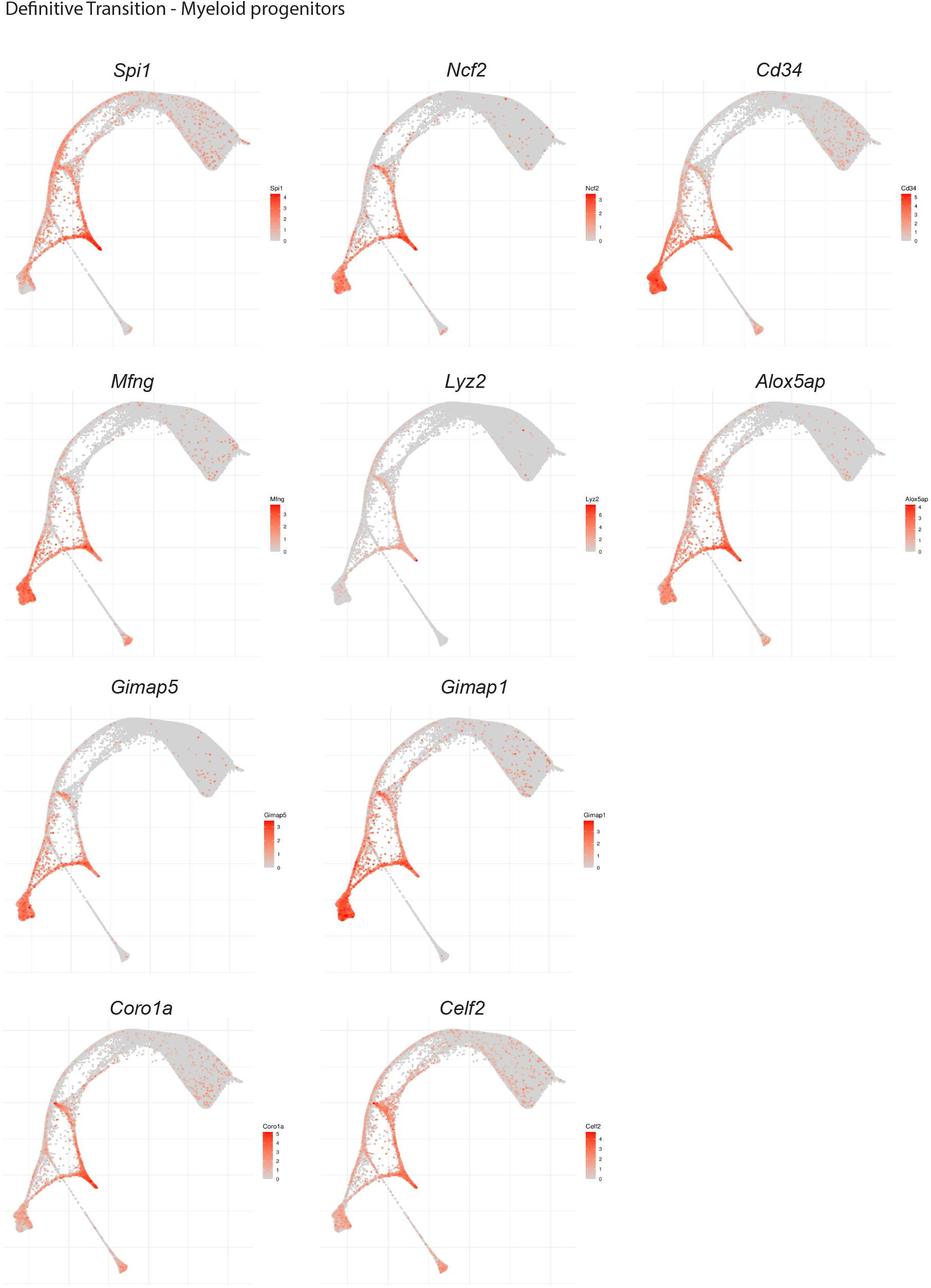
Definitive transition - Myeloid progenitors. Collection of gene markers for definitive blood populations shown as a complement for Figure 2. Force directed layouts displaying gene markers expression levels.

**Supplementary Figure 5.**
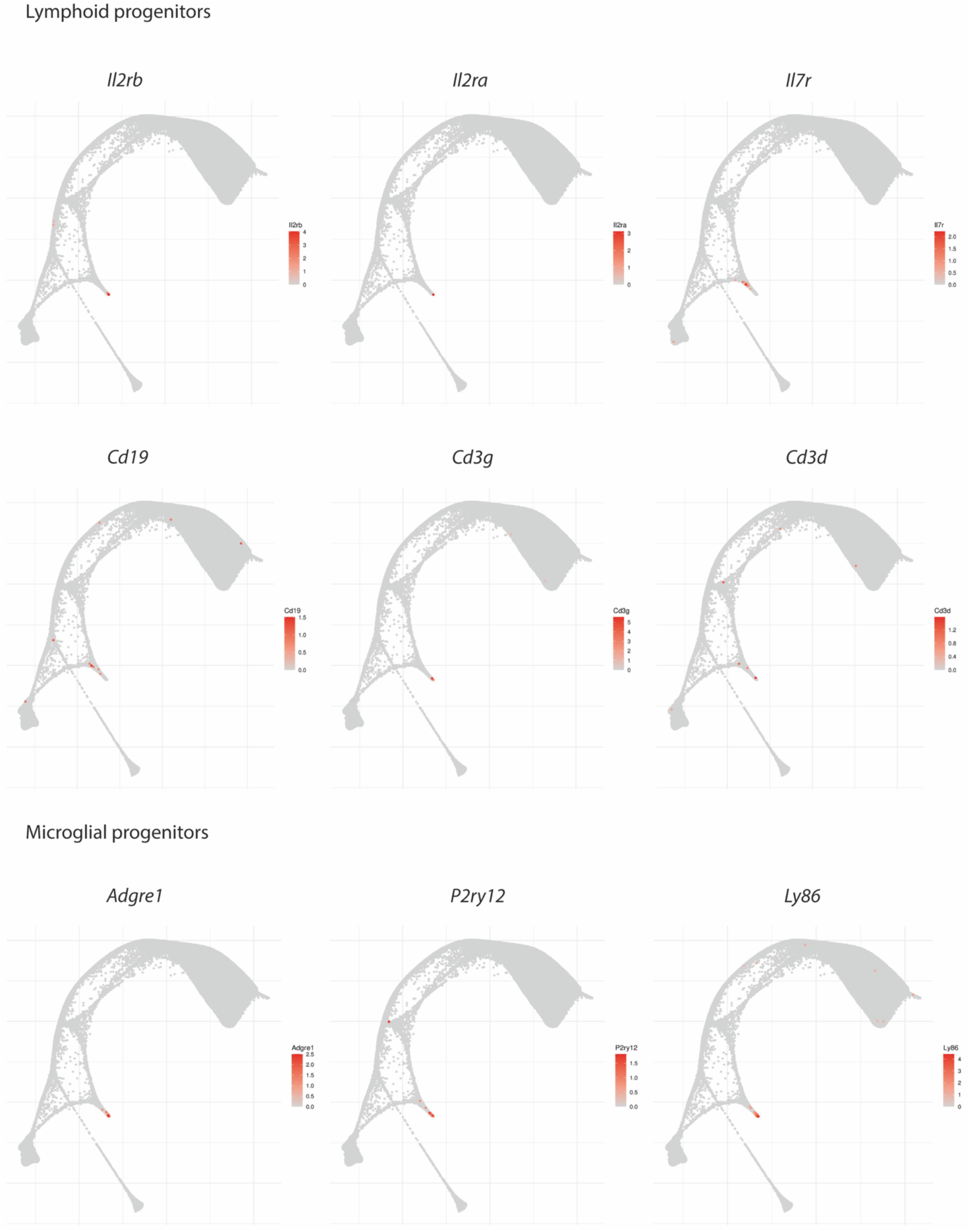
Definitive transition - Lymphoid and microglial progenitors. Collection of gene markers for definitive blood populations shown as a complement for Figure 2. Force directed layouts displaying gene markers expression levels.

**Supplementary Figure 6.**
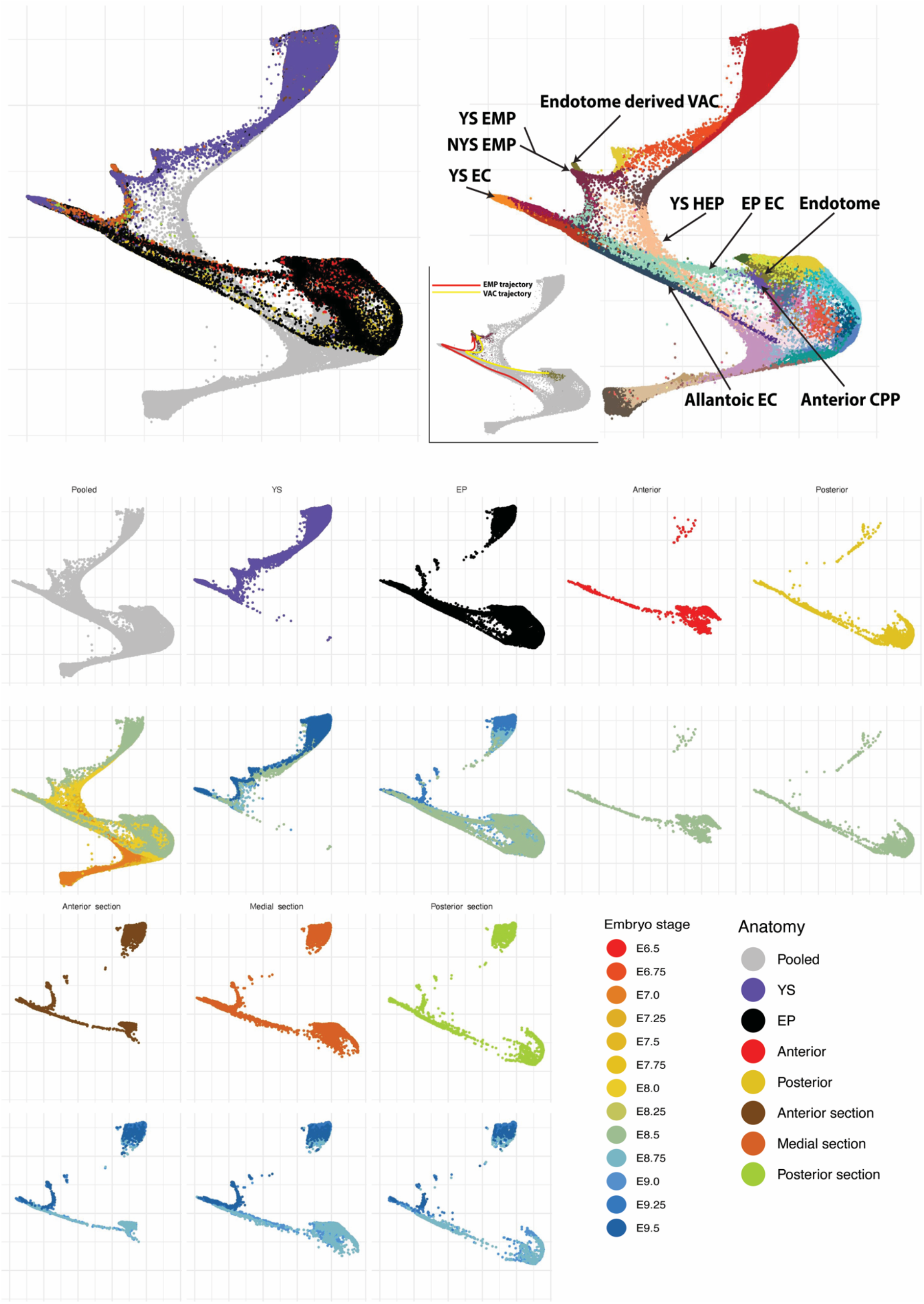
Diverse anatomical origins of the haemato-endothelial landscape. Force directed layout of the haemato-endothelial landscape highlighting anatomical locations and relevant cell type populations (top); each anatomical region is shown in a different panel (bottom). Here, cells are coloured by anatomical locations as well as by embryo stage. Note that embryo proper subdissections where no distinction between anterior, medial and posterior was made before sequencing are labelled as EP.

**Supplementary Figure 7.**
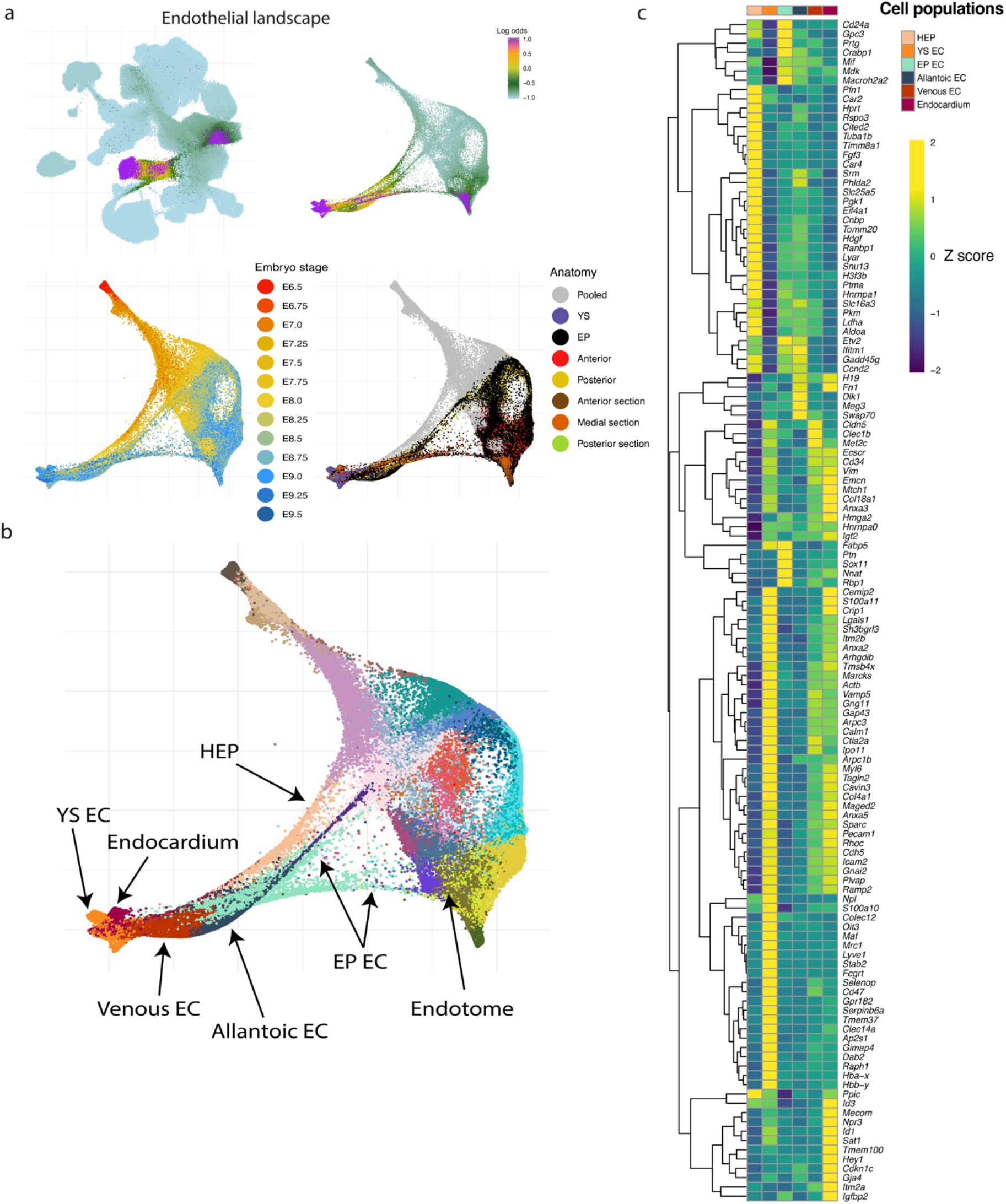
Different origins of endothelial cells. **a)** UMAP layout of the mouse extended atlas displaying the log odds of fate probabilities exclusively associated with endothelial populations (top left). Cells with log odds > - 1 were retained to generate a force directed layout. Cells are coloured by Log odds of fate probabilities of endothelial cells, embryo stage and anatomical region. **b)** Force directed layout of the endothelial landscape highlighting relevant cell populations. HEP: Haemato-endothelial progenitors. EC: Endothelial cell. **c)** Heat map of differentially expressed genes across distinct endothelial populations. Mean gene expression values were computed and scaled by rows (Z-score).

**Supplementary Figure 8.**
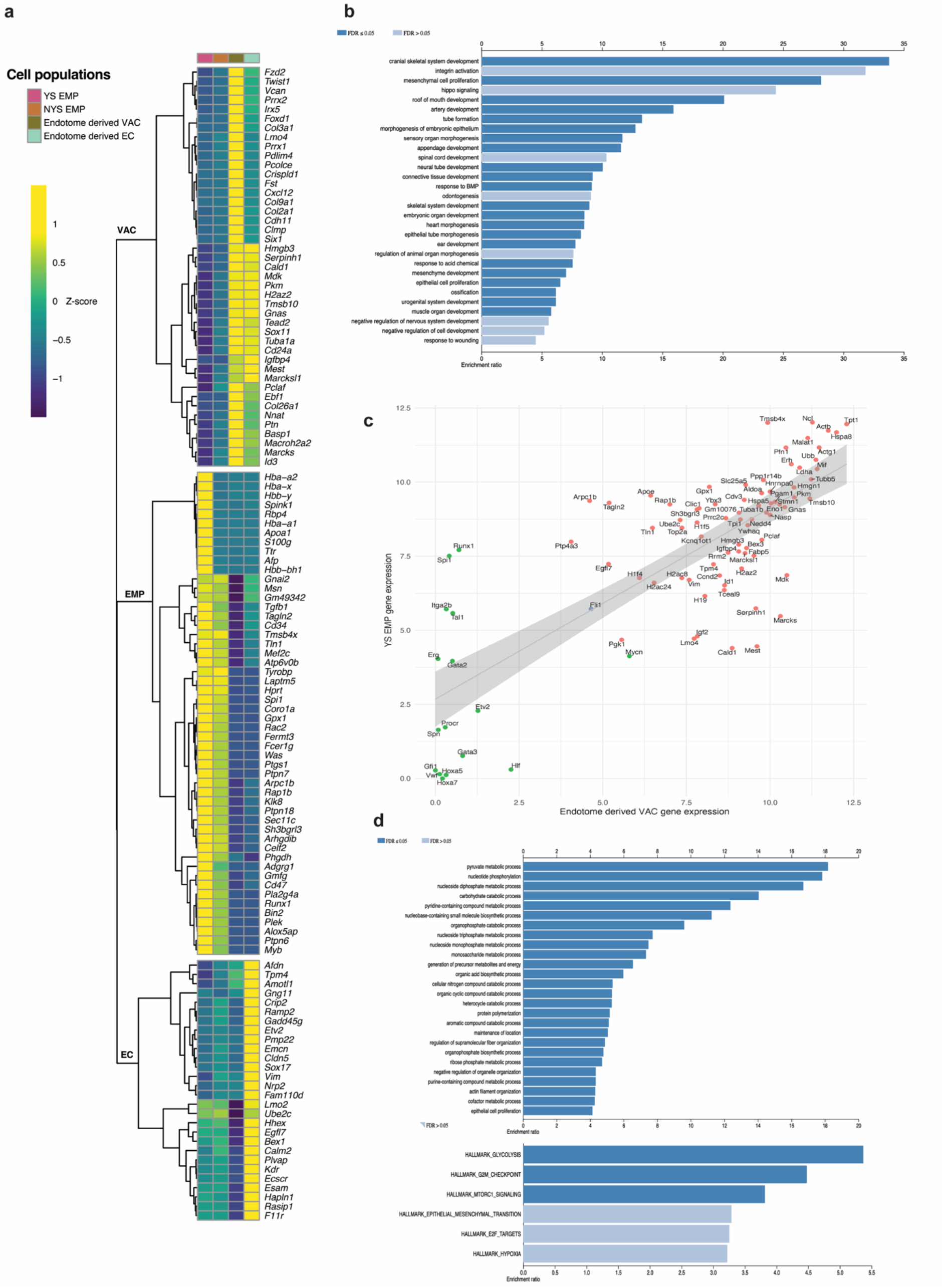
Gene expression signature of endotome populations. **a)** Heat map displaying differentially expressed genes across different cell populations clustered together in the region highlighted in Fig.3. Dii. YS EMP: Yolk Sack EMP, NYS EMP: Non-Yolk Sack EMP, Endotome derived VAC: Endotome derived vascular associated cells (This heat map is an extended version of the analysis performed for Fig.3.E). Mean gene expression values were computed and scaled by rows (Z-score). **b)** Gene set over representation analysis (ORA) performed with *webgestalt*[Liao et al., 2019] against Gene Ontology Biological Processes of VAC genes highlighted in panel A. Significant terms (FDR < 0.1) are generally associated with connective tissue development. **c)** Gene correlation analysis between YS EMPs and endotome derived VACs (red dots: positively correlated genes identified by CCA, green dots: manually selected genes associated with HSC progenitors, blue dots: Intersecting genes between both). Metacells were used to strengthen the gene expression signals (see methods). **d)** ORA of positively correlated genes against hallmark gene sets (FDR < 0.1) and Gene Ontology Biological Processes (FDR < 0.05). The resulting hallmarks reflect cell growth (evidenced by cell cycle genes, mTORC1 signalling and E2f2 targets), glycolytic metabolism and *Myc* activation (the latter above the significance threshold) as well as epithelial-mesenchymal transition while no evidence of haematopoietic identity, but leaving the possibility of niche function akin to the situation observed in Zebrafish.

**Supplementary Figure 9.**
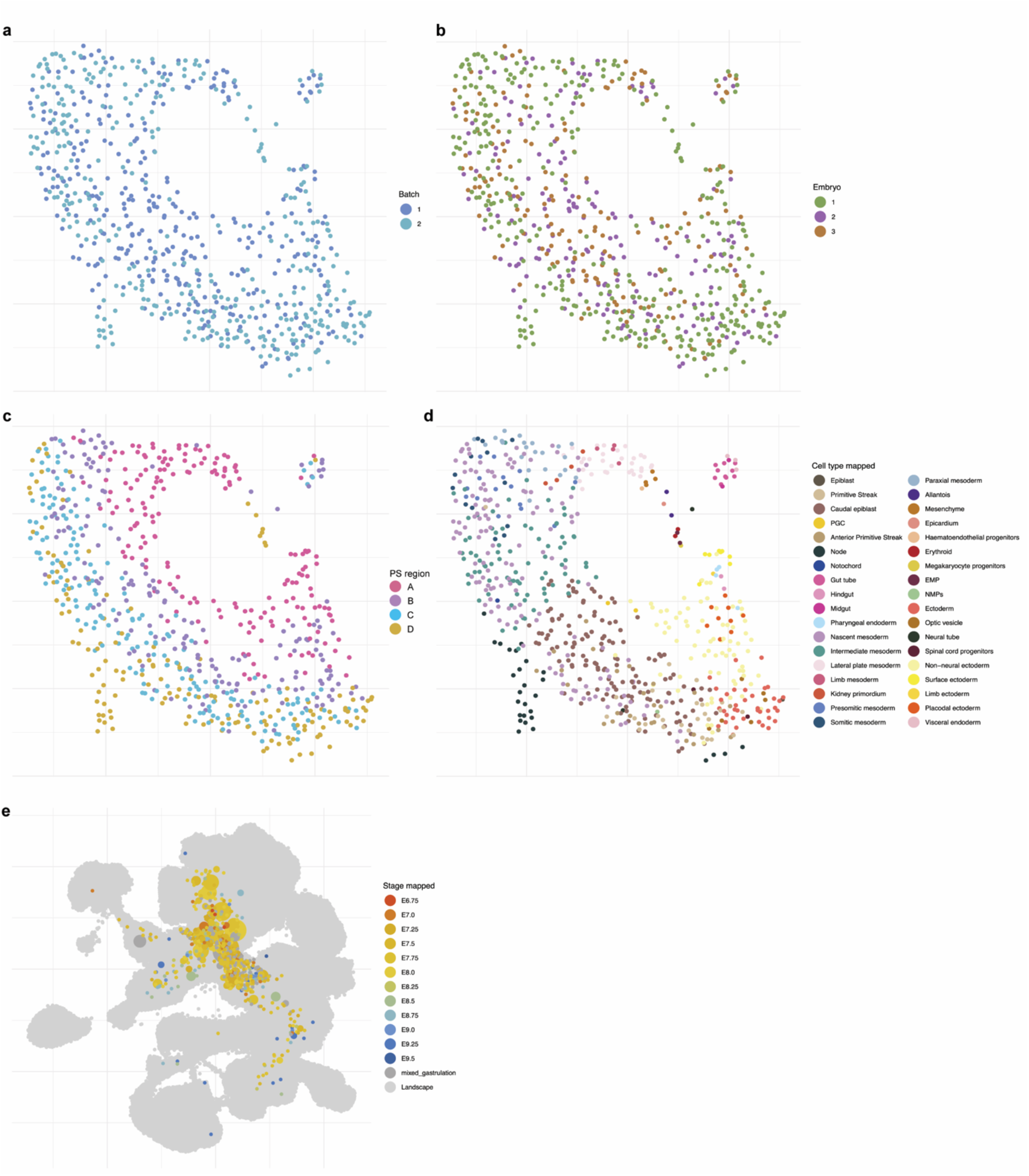
Primitive streak dissections profiled with Smart-seq2. **a)** UMAP layout of single cell transcriptomes from primitive streak dissections after batch correction (cells are coloured by batch). **b)** Three different embryos were profiled (cells are coloured by dissected embryos). **c)** Dissected portions of the Primitive Streak from Anterior to Posterior (cells are coloured by portions). **d)** Cell type annotations assigned to cells by transferring annotation from the whole atlas (cells are coloured by cell types). **e)** UMAP of primitive streak cells mapped onto the extended mouse atlas. Cells are coloured by the embryo stage corresponding to its closest neighbour in the atlas.

**Supplementary Figure 10.**
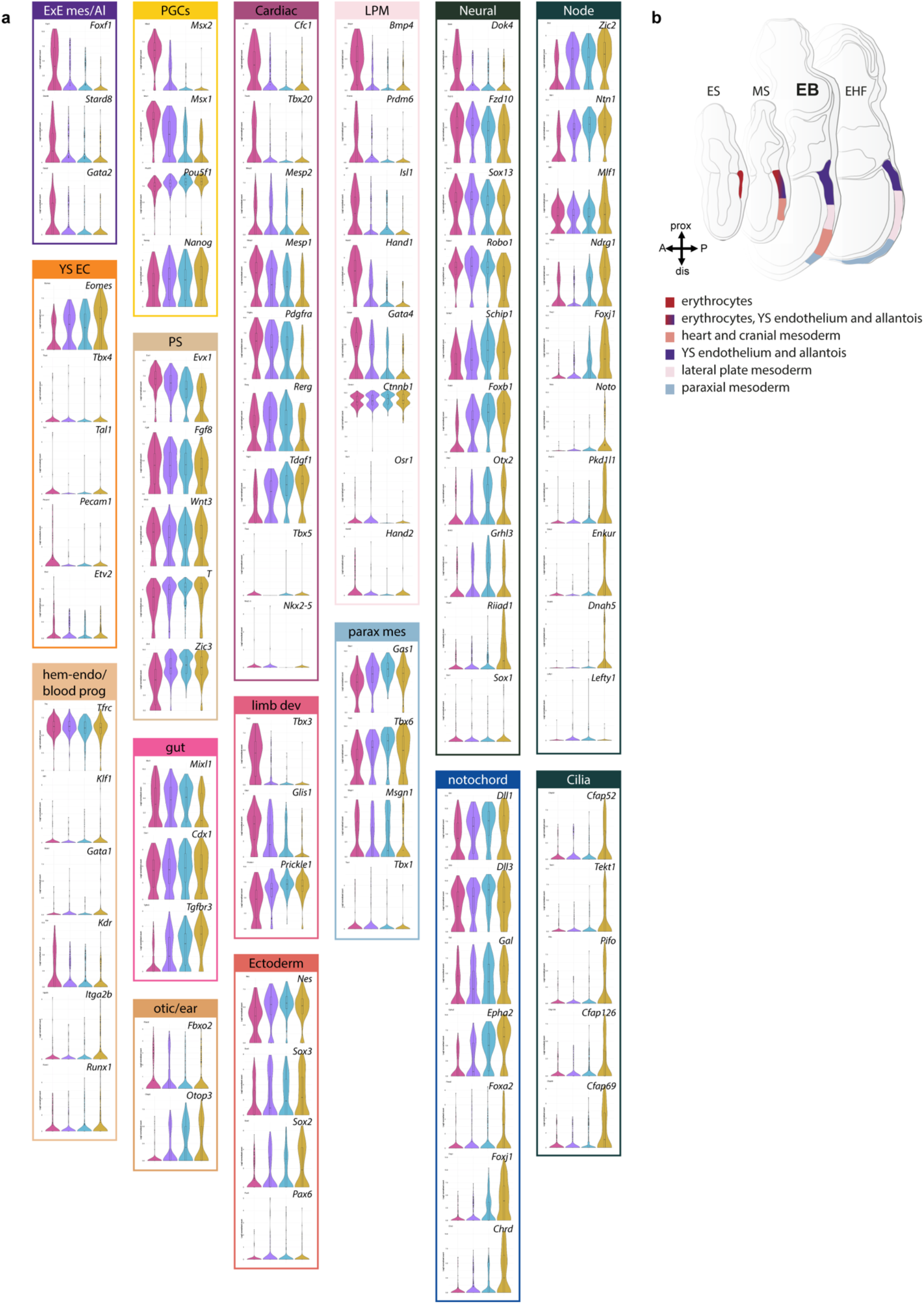
Examples of cell type-specific gene signatures within primitive streak region A to D. **a)** Violin plots of cell type-specific gene signatures within the 4 primitive streak regions are shown. Cell types are colour coded according to the UMAP legend (Fig.4b). **b)** Schematic representation of previously established primitive streak fates at different embryonic stages during gastrulation, adapted from Kinder et al., 2001.

**Supplementary Figure 11.**
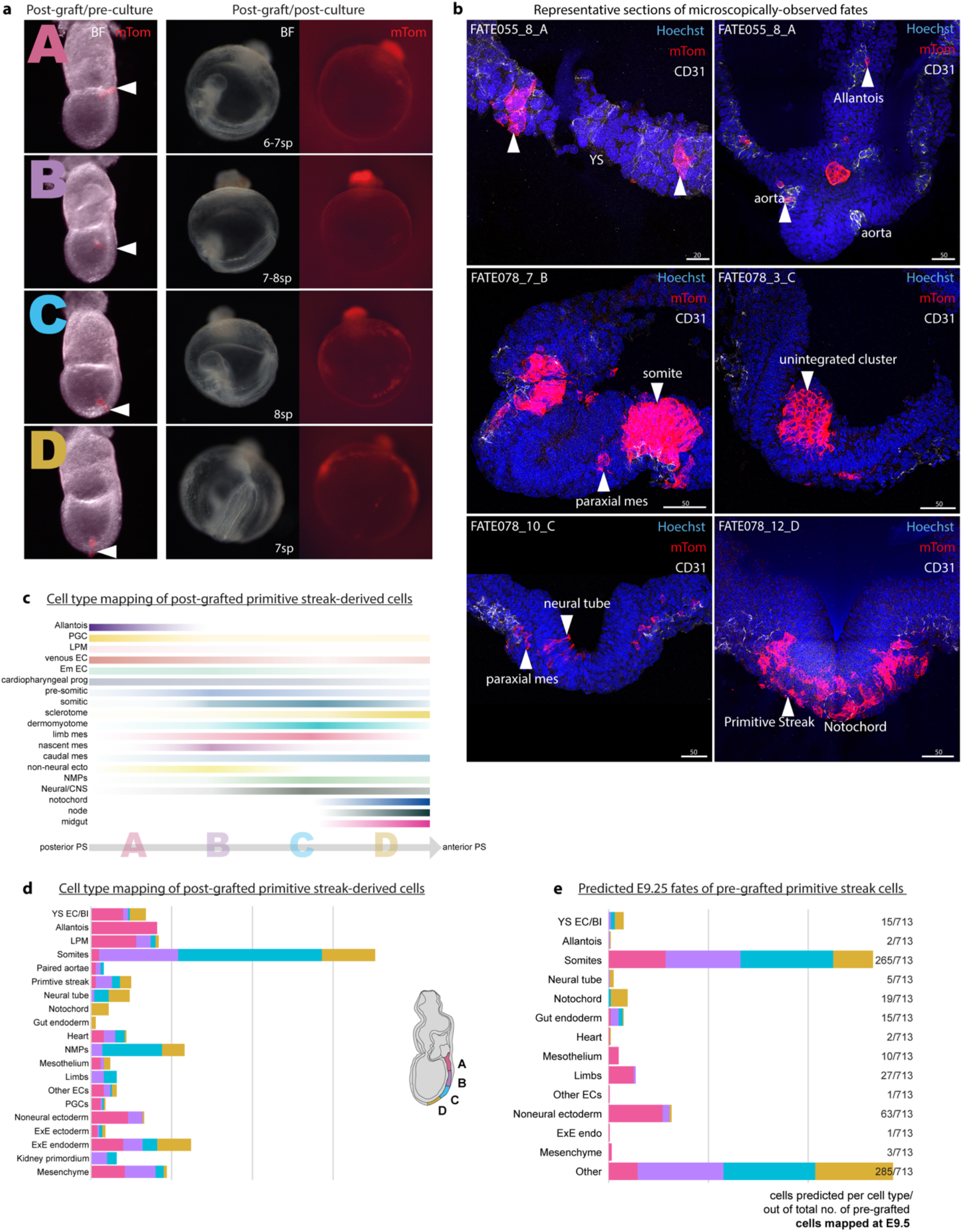
Grafting controls and full fate maps. **a)** Wholemount images showing EB-stage wild type embryos orthotopically grafted with primitive streak regions A-D of mTom transgenic mouse embryos. Arrowheads point to the grafting site along the primitive streak axis with visible red cluster cells lodged in the primitive streak immediately after grafting and before embryo culture. After 24h culture, embryos developed around 7 somite pairs (Suppl.Table 1), had a beating heart and showed mTom contribution in their tissues. **b)** Representative immunofluorescent images of post-grafted/post-cultured embryo sections, where mTom contribution was assessed based on microscopic observations. **c)** Percentile representation of cell type mapping along the posterior-anterior axis of the primitive streak (from A to D, respectively) of post-grafted cells. The colour intensity indicates cell type abundance and resembles the gradient of cell type bias from specific primitive streak regions. **d)** Full fate map of transcriptionally observed fates based on the mapping of the post-grafted cells on the extended atlas. **e)** Full fate map of predicted fates of the pre-graft PS portion cells, determined with the Waddington-OT algorithm. Prediction was made including the full extended atlas, thus up to E9.25.

